# The first complete genome of the extinct European wild ass (*Equus hemionus hydruntinus*)

**DOI:** 10.1101/2023.06.05.543765

**Authors:** Mustafa Özkan, Kanat Gürün, Eren Yüncü, Kıvılcım Başak Vural, Gözde Atağ, Ali Akbaba, Fatma Rabia Fidan, Ekin Sağlıcan, N. Ezgi Altınışık, Dilek Koptekin, Kamilla Pawłowska, Ian Hodder, Sarah E. Adcock, Benjamin S. Arbuckle, Sharon R. Steadman, Gregory McMahon, Yılmaz Selim Erdal, C. Can Bilgin, Eva-Maria Geigl, Anders Götherstrom, Thierry Grange, İnci Togan, Füsun Özer, Mehmet Somel

## Abstract

We present paleogenomes of three morphologically-unidentified Anatolian equids dating to the 1^st^ millennium BCE, sequenced to coverages of 0.6-6.4X. Mitochondrial DNA haplotypes of the Anatolian individuals clustered with those of *Equus hydruntinus* (or *Equus hemionus hydruntinus*), the extinct European wild ass. The Anatolian wild ass whole genome profiles fall outside the genomic diversity of other extant and past Asiatic wild ass (*E.hemionus*) lineages. These observations strongly suggest that the three Anatolian wild asses represent *E.hydruntinus*, making them the latest recorded survivors of this lineage, about a millennium later than the latest observations in the zooarchaeological record. Comparative genomic analyses suggest that *E.hydruntinus* was a sister clade to all ancient and present-day *E.hemionus* lineages, representing an early split. We also find indication of gene flow between hydruntines and Middle Eastern wild asses. Analyses of genome-wide heterozygosity and runs of homozygosity reveal that the Anatolian wild ass population had severely lost genetic diversity by the mid-1^st^ millennium BCE, a likely omen of its eventual demise.

## Introduction

Since its paleontological description more than a century ago (Regalia, 1907), the “European wild ass”, *Equus hydruntinus* (or *Equus hemionus hydruntinus*), has remained an enigmatic creature (Bennett et al., 2017; Boulbes & van Asperen, 2019; Burke et al., 2003; Geigl & Grange, 2012; Orlando et al., 2006). It was a gracile non-caballine equid, once roaming open and dry habitats in Europe and Southwest Asia (Figure 1A) and featuring in Upper Paleolithic cave art (BernÁldez-Sánchez & García-Viñas, 2019; Cleyet-Merle & Madelaine, 1991; Bennett et al., 2017) and on Neolithic pottery (Bennett et al., 2017). Its history in the fossil record starts with the late Middle or Late Pleistocene and ends within the Holocene, when it goes extinct (Boulbes & van Asperen, 2019; Crees & Turvey, 2014; Geigl & Grange, 2012).

**Figure 1.**
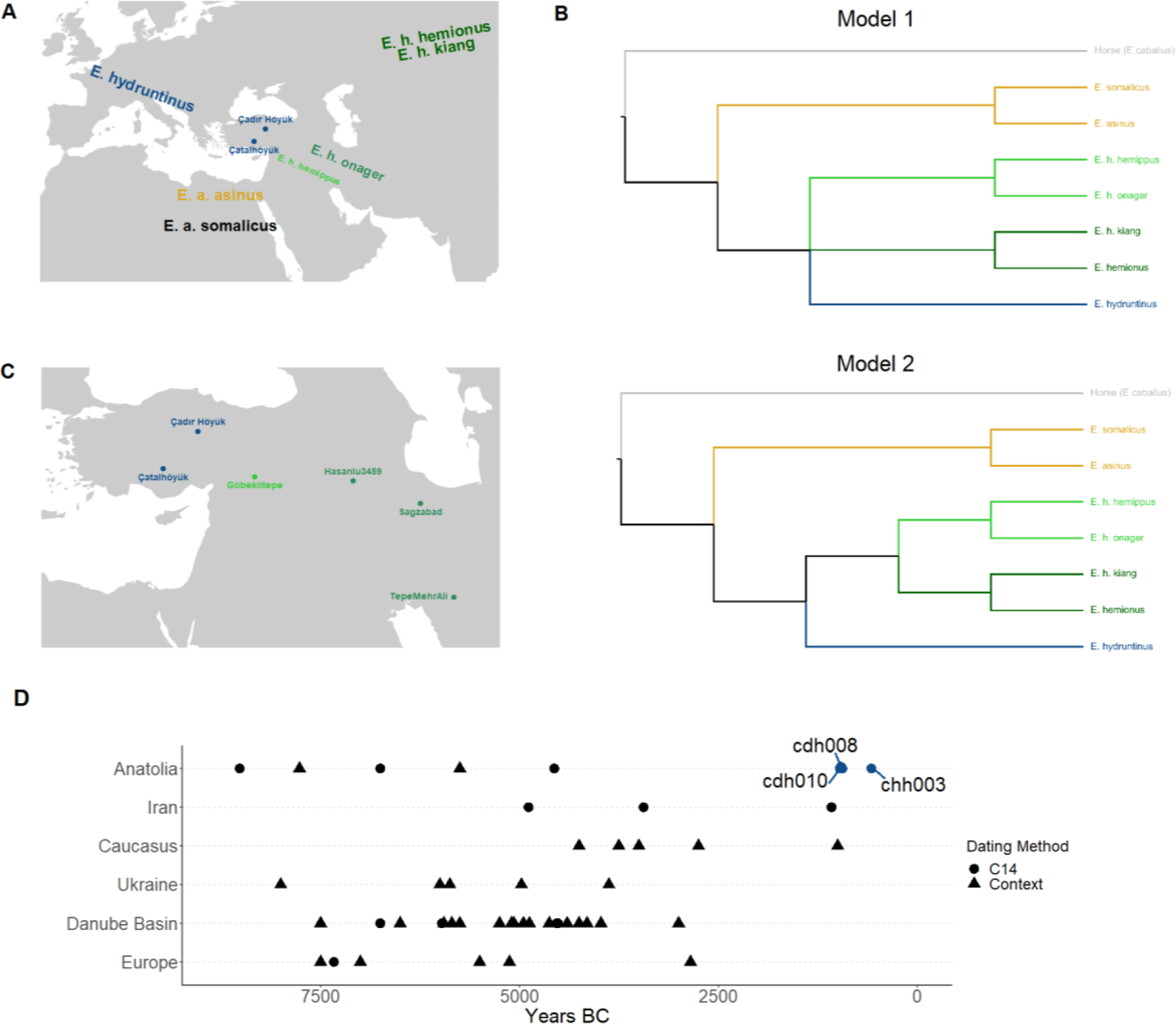
**A)** The map shows the geographical locations of Çadır Höyük and Çatalhöyük (the excavation sites where the wild ass remains analyzed in this study were recovered), as well as the approximate dispersal areas of extant and extinct ass taxa. **B)** Models summarizing two hypotheses regarding the taxonomic position of European wild ass. **C)** Also shows the locations of three ancient hemione individuals and one hemippe individual the partial genomes of which were previously published (Fages et al., 2019; Bennett et al., 2022). **D)** Timeline of dated *E. hydruntinus* reportings with samples reported in this study (adapted after Crees and Turvey, 2014).

Early research in the 20^th^ century comparing *E. hydruntinus* remains with those of other equids identified resemblances to diverse taxa, including the African asses (*E. asinus*) and the zebra (*E. zebra*), the Asiatic asses (*E. hemionus*), or the extinct stenonine horses, leaving its phylogenetic position disputed for many decades (Azzaroli, 1991; Davis, 1980; Forsten & Ziegler, 1995; Stehlin & Graziosi, 1935; Eisenmann & Baclay, 2000; Eisenmann & Mashkour, 2000). Within the last two decades, osteological studies on new fossil findings concluded that *E. hydruntinus* was systematically closer to extant Asian asses, *i.e.* hemiones, than to African asses or zebras (Burke et al., 2003; Eisenmann & Mashkour, 1999; Orlando et al., 2006) (Figure 1A). Ancient DNA analyses of mitochondrial DNA (mtDNA) supported this conclusion: partial and complete mtDNA sequences from *E. hydruntinus* and extant hemiones (the kiang of Tibet; the kulan of Mongolia; the kulan of Turkmenistan; the onager of Iran) were found to cluster together to the exclusion of other equids (Bennett et al., 2017; Catalano et al., 2020; Orlando et al., 2006, 2009). Moreover, these mitochondrial DNA studies suggested that the hemione - hydruntine division may represent taxonomic over-splitting (Bennett et al., 2017; Orlando et al., 2006, 2009). Bennett et al. (2017) pointed out that in their mtDNA analyses, hemiones and hydruntines did not appear reciprocally monophyletic. This could be explained by rapid diversification of the *E. hemionus* lineages, including hydruntines, creating an unresolved radiation node (Model 1 in Figure 1B). Accordingly, hydruntines could also be considered a subspecies of *E. hemionus*, *E.h.hydruntinus*, similar to the kulan (*E.h.kulan*) and onager (*E.h.onager*), considered *E. hemionus* subspecies by the IUCN (Kaczensky et al., 2015).

This suggestion, however, has been contentious. The phylogenetic patterns described were only based on partial mtDNA sequences and it remained possible that analyses of full mtDNA and of nuclear loci could reveal different patterns, *e.g.* an early hydruntine-hemione split (Model 2 in Figure 1B). Supporting an early-split idea, osteological analyses have suggested that *E. hydruntinus* carried a number of unique adaptations distinct from other hemiones, such as a short and wide muzzle adapted to its cold and dry conditions (Boulbes & van Asperen, 2019; van Asperen, 2012). Given the equivocal evidence, there have been calls for in-depth morphometric analyses (Twiss et al., 2017) and for the analysis of nuclear genomic data to resolve the issue (Boulbes& van Asperen, 2019; Crees & Turvey, 2014).

Another controversy surrounding *E. hydruntinus* or *E.h.hydruntinus,* which hereinafter we will name hydruntine, involves its extinction dynamics. Crees and Turvey (2014) studied Holocene zooarchaeological records of hydruntines along with paleovegetation data, suggesting that during the Holocene, the hydruntine range was highly fragmented and restricted to regions with relatively open habitats, such as the Danube basin and the Anatolian steppe. The authors predicted its extinction in the Danube region by the third millennium BCE, and in Iran and South Caucasus possibly within the first millennium BCE. However, other scholars have suggested hydruntines in Iran and in Anatolia could have gone extinct by the 2^nd^ millennium BCE (Mashkour et al., 1999; Guimaraes et al., 2020). Its extinction is generally attributed to a combination of factors including increased aridity associated with the 4.2-ka event, competition with livestock for pasture resources, and hunting. Nevertheless, due to the relative rarity of hydruntines in the zooarchaeological record (compared to *e.g.* red deer) and also due to difficulties in morphological identification (Geigl & Grange, 2012; Twiss et al., 2017), the timing of hydruntine extinction remained largely uncertain (Boulbes & van Asperen, 2019; Crees & Turvey, 2014; Nores et al., 2015).

Here we present the first full genomic data genetically attributable to the hydruntine, obtained from three Anatolian equids from 1^st^ millennium BCE Central Anatolia. The analysis of these genomes allows us to resolve questions on the phylogenetics, demographic history and extinction dynamics of this taxon.

## Results

We performed ancient DNA extraction on 15 equid tooth and bone samples excavated from two Central Anatolian sites, 11 from Çatalhöyük and four from Çadır Höyük (Figure 1C). We generated genome-wide genetic data from these samples using shotgun sequencing (Supplementary Table 1) and obtained 0.02%-11.35% (median=0.07%) of endogenous DNA by mapping to the horse reference genome (version EquCab2.0). We chose three samples (cdh008, cdh010 and chh003) with >5% endogenous DNA all of which exhibited typical postmortem damage (PMD) signals for further sequencing (Table 1, Supplementary Figure 1). We thus produced three genomes with nuclear coverages 6.38x, 0.72x and 0.57x (Table 1, Supplementary Table 2). Radiocarbon dating of the skeletal material placed all three individuals within the early/mid 1^st^ millennium (Iron Age) in Anatolia (Table 1, Figure 1D). We also generated a near-complete mitogenome sequence of the highest coverage (140.04x) sample cdh008 (Methods).

**Table 1.**
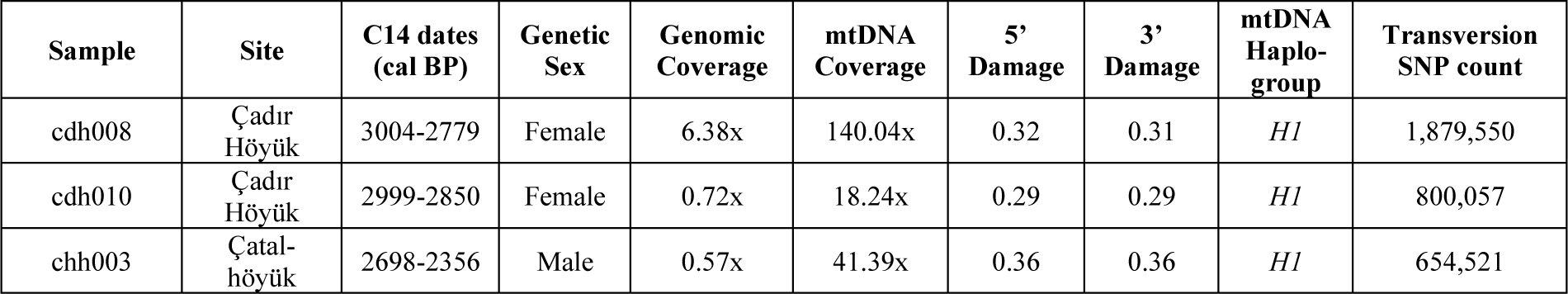
Genome information and radiocarbon dates of the three newly sequenced ancient Anatolian equids. Genome coverages are provided as average read depth across all positions on the whole genome or the mitochondrial genome, after removing duplicate reads. “5’ Damage” and “3’ Damage” represent postmortem damage-induced substitution frequencies (C->T and G->A), given as fractions of C or G nucleotides at the 5’ and 3’ ends, respectively. “SNP count” refers to SNPs called among a total of 2,146,416 variable autosomal transversion positions identified among 12 *E.asinus* and *E.heminonus* genomes (Methods). Radiocarbon (C14) date intervals (95%), calibrated using IntCal20 (Reimer et al., 2020) curve, are given as calibrated years Before Present (BP). The raw radiocarbon dates and laboratory codes are available in Archaeological Supplement Table 2.

| Sample | Site | C14 dates (cal BP) | Genetic Sex | Genomic Coverage | mtDNA Coverage | 5’ Damage | 3’ Damage | mtDNA Haplo-group | Transversion SNP count |
| --- | --- | --- | --- | --- | --- | --- | --- | --- | --- |
| cdh008 | Çadır Höyük | 3004-2779 | Female | 6.38x | 140.04x | 0.32 | 0.31 | H1 | 1,879,550 |
| cdh010 | Çadır Höyük | 2999-2850 | Female | 0.72x | 18.24x | 0.29 | 0.29 | H1 | 800,057 |
| chh003 | Çatalhöyük | 2698-2356 | Male | 0.57x | 41.39x | 0.36 | 0.36 | H1 | 654,521 |

### Genetic analyses assign the Iron Age Anatolian wild ass individuals to E.h.hydruntinus/*E. hydruntinus*

During the Bronze and Iron Ages, horses and the domestic donkey were common in Anatolia, whilst hemiones were present in southern-eastern Anatolia, and hydruntine populations might also have survived in the region (Archaeological Supplementary File Table 1, see references). Due to a lack of sufficient diagnostic characteristics, the three specimens were each identified to the level of the genus *Equus* but no attempt was made to assign species based on morphological criteria (Archaeological Supplementary File Table 2). As a first step towards characterizing the three Anatolian individuals, we used the *Zonkey* pipeline, which performs taxonomic classification of equid genomic data by comparing them with a reference genome panel (Schubert et al., 2017). *Zonkey* classified all three individuals as “Asian wild ass-related” (Supplementary Table 1).

We then compared mtDNA data from these three Anatolian equids with published partial mtDNA sequences. In total, sequences from 82 hemiones (Bennett et al., 2017; Huang et al., 2015; Orlando et al., 2009; Wang et al., 2020) and 21 morphologically-identified hydruntine individuals (Bennett et al., 2017; Catalano et al., 2020; Orlando et al., 2009; Schubert et al., 2016) as well as 7 other equids (Bennett et al., 2017; Jónsson et al., 2014; Wang et al., 2020) were collected. We used a 249 bp-long fragment, the longest common fragment across all available samples, to construct a haplogroup network; we also used a 361 bp-long D-loop fragment for *BEAST* analysis (Supplementary Table 3). Both in a haplogroup network and in Bayesian trees, mtDNA sequences of the three Anatolian equids clustered with published hemione-hydruntine sequences with strong support (Supplementary Figures 2 and 3). Moreover, all three new sequences were a sub-branch of the *H1* haplotype clade [as defined by (Bennett et al., 2017)]. *H1* is the most prevalent mtDNA haplotype among the 19 hydruntine individuals in this dataset, appears exclusive to hydruntine, and was already detected among Anatolian hydruntines of Neolithic and Bronze Ages, c.7,850-2,500 BCE (Bennett et al., 2017) (Supplementary Figure 2). Together, these results indicate that the mitochondrial lineage of the three Anatolian equids belonged to wild asses, and was more closely related to hydruntines than to hemiones.

We investigated this further using complete mitochondrial DNA sequences, taking advantage of a recently published almost complete mtDNA sequence of a morphologically-identified *E.h.hydruntinus/E.hydruntinus* specimen (Catalano et al., 2020). We compiled full mtDNA sequences from a total of 9 ancient and modern-day equids and outgroup lineages (Methods, Supplementary Table 3). Again, both Bayesian trees constructed using *BEAST* (Figure 2A), as well as neighbour-joining and maximum likelihood trees (Supplementary Figure 4) clustered the Anatolian wild asses with the published hydruntine with full support.

**Figure 2.**
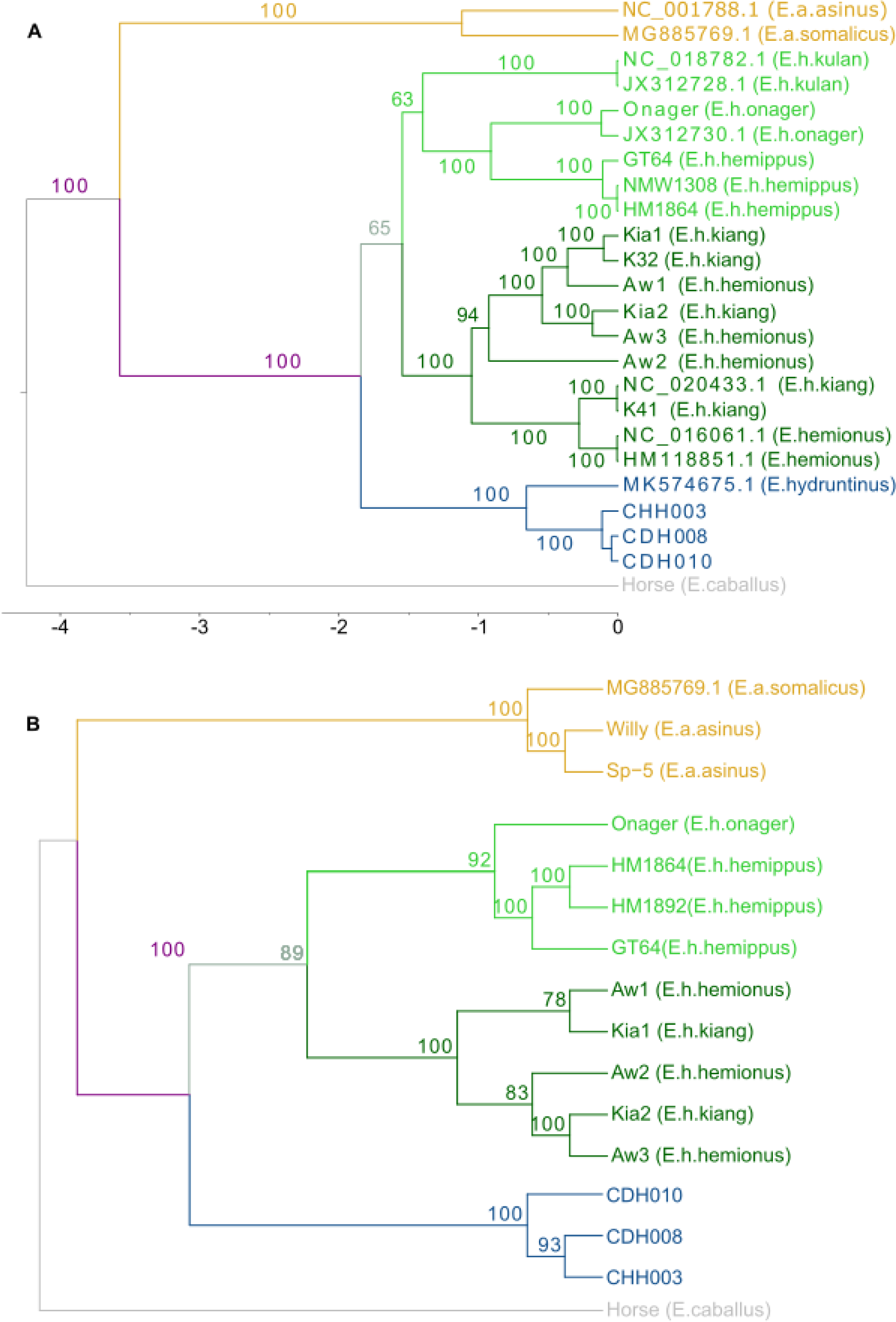
**A)** A Bayesian phylogenetic tree constructed using a 13,042 bp long consensus sequence of the whole mitochondrial DNA among various equid lineages. Numbers on the internal nodes show posterior probability support. **B)** A maximum likelihood tree constructed using concatenated protein coding sequences (29,384,180 bp) across the 15 genomes using only transversion sites. Numbers on the internal nodes show bootstrap support. External nodes indicate the genome/individual IDs, with species names in parentheses. A version showing branch lengths can be found in Supplementary Figure 5.

Mitochondrial DNA phylogenies may be incongruent with the overall phylogenetic history of a species due to drift, selection or incomplete lineage sorting (Toews & Brelsford, 2012). We hence explored the whole genome data for evidence that the three Anatolian wild asses might represent a hydruntine. For this, we compiled a list of c.2 million autosomal SNPs positions using published asinus and hemionus genomes (Methods). We genotyped the three Anatolian wild ass genomes, as well as 13 published modern-day ass and donkey genomes at these positions (Table 1, Supplementary Table 3). For comparison, we further included three published ancient onager genomes from Iran (TepeMehrAli, Hasanlu3459 and Sagzabad) (Fages et al., 2019), and three ancient hemippe genomes, one from Göbeklitepe, Turkey (GT64), and two from museum specimens (Hm_1864 and Hm_1892) (Bennett et al., 2022) (Figure 1C). We then constructed a multidimensional scaling (MDS) summary plot of genomic diversity based on the outgroup *f^3^*-statistic (Methods), including the modern-day African and Asian asses and the ancient individuals (Figure 3; Supplementary Figure 6, Supplementary Table 4). This revealed a unique position for the three Anatolian wild asses, distinct from African asses as well as the Asian hemiones, including ancient onagers from Iran and the hemippes. Accordingly, the three Anatolian wild asses appear phylogenetically distinct from either the hemione or the hemippe clades. The hydruntine is the only other wild ass lineage identified in the fossil record to have lived in Holocene Anatolia. Together with the mtDNA evidence, this strongly suggests the three wild asses belonged to the extinct hydruntine lineage.

**Figure 3.**
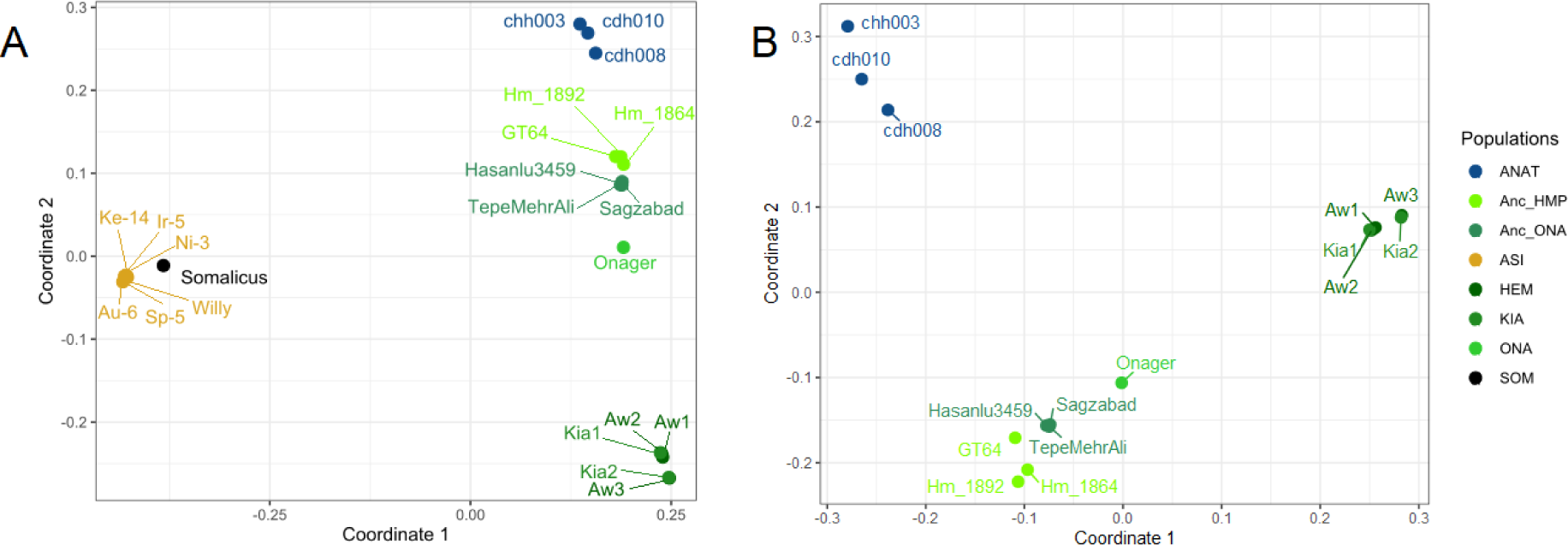
Multidimensional scaling plot of A) donkey and wild asses and B) only wild asses generated using a distance matrix based on (1-outgroup *f*^3^) statistics calculated from the autosomal variation dataset consisting of 2,146,461 sites (ANAT: Anatolian samples reported in this article; Anc_HMP: Ancient *E.h.hemippus*; Anc_ONA: Ancient *E.h.onager*; ASI: *E.asinus*; HEM: *E.hemionus*; KIA: *E.h.kiang*; ONA: *E.h.onager*; SOM: *E.somalicus*).

### Whole genome analyses support hemiones and hydruntines as sister clades

Because the hydruntine has yet only been genetically described through mtDNA, we leveraged upon the new Anatolian genomes to investigate evolutionary relationships among asses. Based on partial mtDNA sequences, it had been previously suggested that hydruntines were most closely related to hemiones, to the exclusion of other equids (Bennett et al., 2017; Catalano et al., 2020; Orlando et al., 2006, 2009). Our results above are fully consistent with this hypothesis, with Anatolian and modern-day Asian wild ass individuals clustering to the exclusion of African asses in mtDNA analyses (Figure 2, Supplementary Figures 3 and 4). Anatolian wild asses are also closer to hemiones in the MDS- and heatmap-based summaries of nuclear genomic variation using outgroup-*f*^3^ scores (Figure 3; Supplementary Figure 6 and 7, Supplementary Table 4). We further tested this pattern with D-statistics, employing the c.2 million autosomal transversion SNP panel (Methods) and using the horse as an outgroup (Supplementary Table 5 and 6). D-tests of the form *D(Horse, Anatolia; Africa, Hemione)* and *D(Horse, Hemione; Africa, Anatolia)* were all significantly positive (*Z*>3; Methods), in line with the notion that hemiones and hydruntines represent sister taxa (Figure 4; Supplementary Tables 4 and 5). We replicated the same results using c.2 million transversions in a secondary variation panel (Schubert et al., 2017) (Methods) (Supplementary Figure 9).

**Figure 4.**
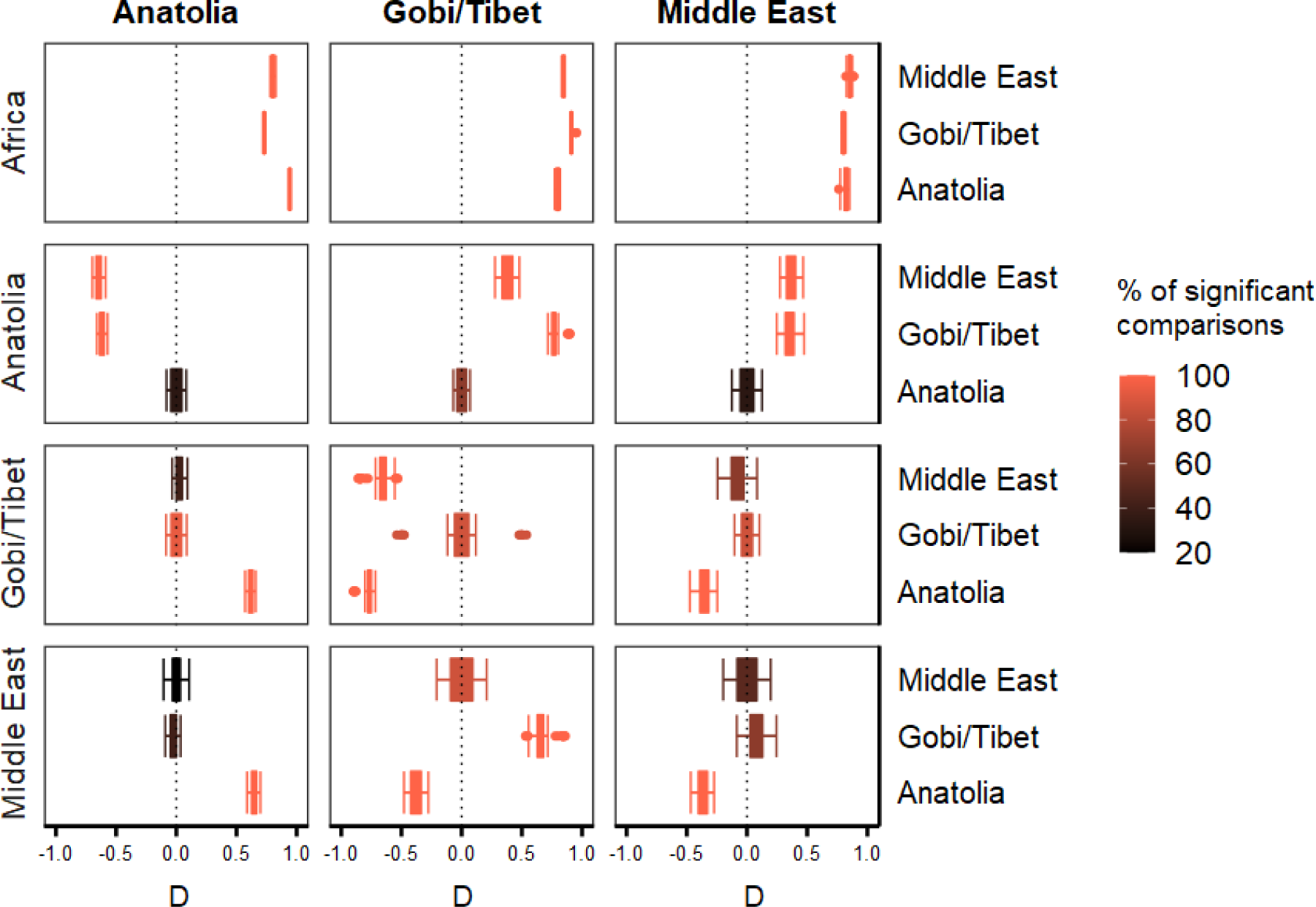
D-statistics calculated using autosomal data between regional groups. Test groups are shown on the top. The color gradient from pink to black represents the fraction of comparisons that are nominally significant (Z>3). Regional groups include: Anatolia (cdh008, cdh010, chh003), Gobi/Tibet (Aw1, Aw2, Aw3, Kia1, Kia2), Middle East (GT64, Hm_1864, Onager, Hasanlu3459, Sagzabad) and Africa (Asinus, Somalicus).

### Hemiones and hydruntines are reciprocally monophyletic

Earlier analyses of partial mtDNA sequences had suggested that the hemione-hydruntine divide could be artificial, with some hemiones appearing closer to hydruntines than to other hemiones (Bennett et al., 2017; Orlando et al., 2009). Meanwhile, the external positioning of hydruntines to hemiones in MDS analyses (Figure 3; Supplementary Figure 6) implied the contrary. We tested this question further through a number of approaches. First, we studied mtDNA trees produced using either partial or full mtDNA sequences. A Bayesian tree based on full sequences included a hydruntine branch external to hemiones with full support (Figure 2A), while a tree with a larger number of samples but based on a shorter fragment (361 bp) had similar topology albeit without statistically significant support (Supplementary Figure 3).

We repeated this phylogenetic analysis using whole genome data of 13 equids and using concatenated protein-coding sequences (Methods). A maximum likelihood tree constructed with this dataset indicated reciprocal monophyly of the Asiatic wild asses (i.e. the 9 Asian wild ass genomes representing the kiang, hemione, onager, and hemippe) and the hydruntine clade (represented by the Anatolian wild asses) (Figure 2B).

We further used D-tests to further clarify hemione-hydruntine relationships. We started by testing monophyly of the Anatolian wild ass genomes with *D(Horse, AnatoliaX; AnatoliaY, Non-Anatolia)*, where *AnatoliaX* and *AnatoliaY* refer to different Anatolian wild ass genomes, and *Non-Anatolia* refers to any individual hemione, kiang, onager or hemippe genome (Methods; Supplementary Table 5). The three Anatolian wild asses formed their own clade to the exclusion of all Asiatic wild asses, with all 60 relevant comparisons being significant in this direction (*Z*>3; Methods) (Figure 4, Supplementary Table 5). We also conducted reciprocal tests of the form *D(Horse, Non-AnatoliaZ; AnatoliaX, Non-AnatoliaW)*, where *AnatoliaX* refers to any Anatolian wild ass genome, while *Non-AnatoliaZ* and *Non-AnatoliaW* refer to genomes of different hemione, kiang, onager or hemippe individuals. In all comparisons, each hemione individual (modern-day or ancient) showed higher affinity to other hemiones over Anatolian individuals (*Z*>3) (Figure 4, Supplementary Table 5). The evidence thus supports reciprocal monophyly between hemiones/hemippes and hydruntines, represented by Anatolian wild asses.

### Evidence for gene flow between hydruntines and Middle Eastern wild asses

We next asked whether the Anatolian wild asses might be symmetrically related to all Asian wild ass lineages. Following our earlier observations (Figures 2 and 3), here we divided Asiatic wild asses into two groups: hemione and kiangs forming a Gobi/Tibet cluster while onager and hemippe forming the Middle East cluster. We then performed D-tests of the form *D(Horse, Anatolia; MiddleEast, Gobi/Tibet)*, where *MiddleEast* represents onager and hemippe individuals while *Gobi/Tibet* consists of hemione and kiangs. We found affinities of all three Anatolian wild asses towards the modern-day Iranian onager and a museum specimen of hemippe (Hm_1864) over other Asian wild asses, including other onagers and hemippes (27 of 30 comparisons with *Z*>0; 23 of 30 comparisons significant at *Z*>3) (Figure 4; Supplementary Figure 9; Supplementary Table 5). This observation is even more pronounced (100% of 75 comparisons at *Z*>3) in comparisons using the secondary variation panel (Supplementary Figure 10; Supplementary Table 6). This would be compatible with gene flow between onager/hemippe and hydruntine populations in Southwest Asia.

We further investigated this by testing *D(Horse, Anatolia; MiddleEastX, MiddleEastY)* where we compared genetic affinity of Anatolian hydruntines between the onager and/or hemippe individuals. Using the main variation panel we observed slight affinity toward the 19^th^ century hemippe over the other hemippe and onagers (12/12 comparisons with *Z*>0 and 5 with *Z*>3; Supplementary Figure 11). Using the second variation panel we found higher affinity to the modern-day onager over all other Middle East wild ass genomes (12 of 12 comparisons with *Z*>3; Supplementary Figure 12). Thus, although we cannot yet fully determine the exact nature of the possible admixture, we hypothesize that gene flow from hydruntine to the onager and hemippe lineages could explain the observed patterns (see Discussion).

### The timing of the hydruntine and Asian wild ass split

The above analyses suggest that hydruntines split from other Eurasian wild asses early in their history, although they might have admixed with Middle Eastern wild asses lineages in more recent times as reflected in the autosomal data. Hence we can estimate the hydruntine-Asian wild ass split time using mtDNA and also autosomal comparisons between hydruntines and Gobi/Tibet wild assess. We first used *BEAST* for mtDNA split time estimation using the caballine and non-caballine divergence times of 4.25 mya as an anchor (Vilstrup et al., 2013) for calibrating splits (Figure 2A). The mitochondrial divergence estimate between European-Anatolian clade and the Asiatic hemiones was 1.84 [0.38-3.23] mya, while that between the hydruntine specimen from Sicily and Anatolian wild asses was 734 [54-2,056] kya. We note that the split times estimated here tend to be higher than earlier estimates (Vilstrup et al., 2013; Bennett et al., 2017; Jonsonn et al., 2014) but also have wide confidence intervals that include those earlier estimates.

We then used the F(A|B) method for estimating the split time between hydruntines and kiang from the Gobi/Tibet region using autosomal DNA. This method is designed for split time estimation when a high-quality and a low-quality genome are available for the investigated lineages (Green et al. 2010; Mualim et al., 2020). We calculated F(A|B), the ratio of derived alleles in the Anatolian wild ass at sites heterozygous in the kiang genome, as 0.145 (and 0.149 using transversions only). Then, we computed the expected F(A|B) values for different divergence times, using both the theoretical estimate (Mualim et al., 2020) and also via simulations (Baumdicker et al., 2022), and using a range of effective population size estimates for the kiang (Methods). The intersection of the observed and expected statistics suggests a most likely population split between 800 kya and 1 mya (Figure 5). This is younger than our mtDNA split time point estimate but within its confidence interval. This autosomal split time would also be compatible with the first fossil records of *E. hydruntinus* in Europe c.600 kya (Boulbes and van Asperen, 2019).

**Figure 5.**
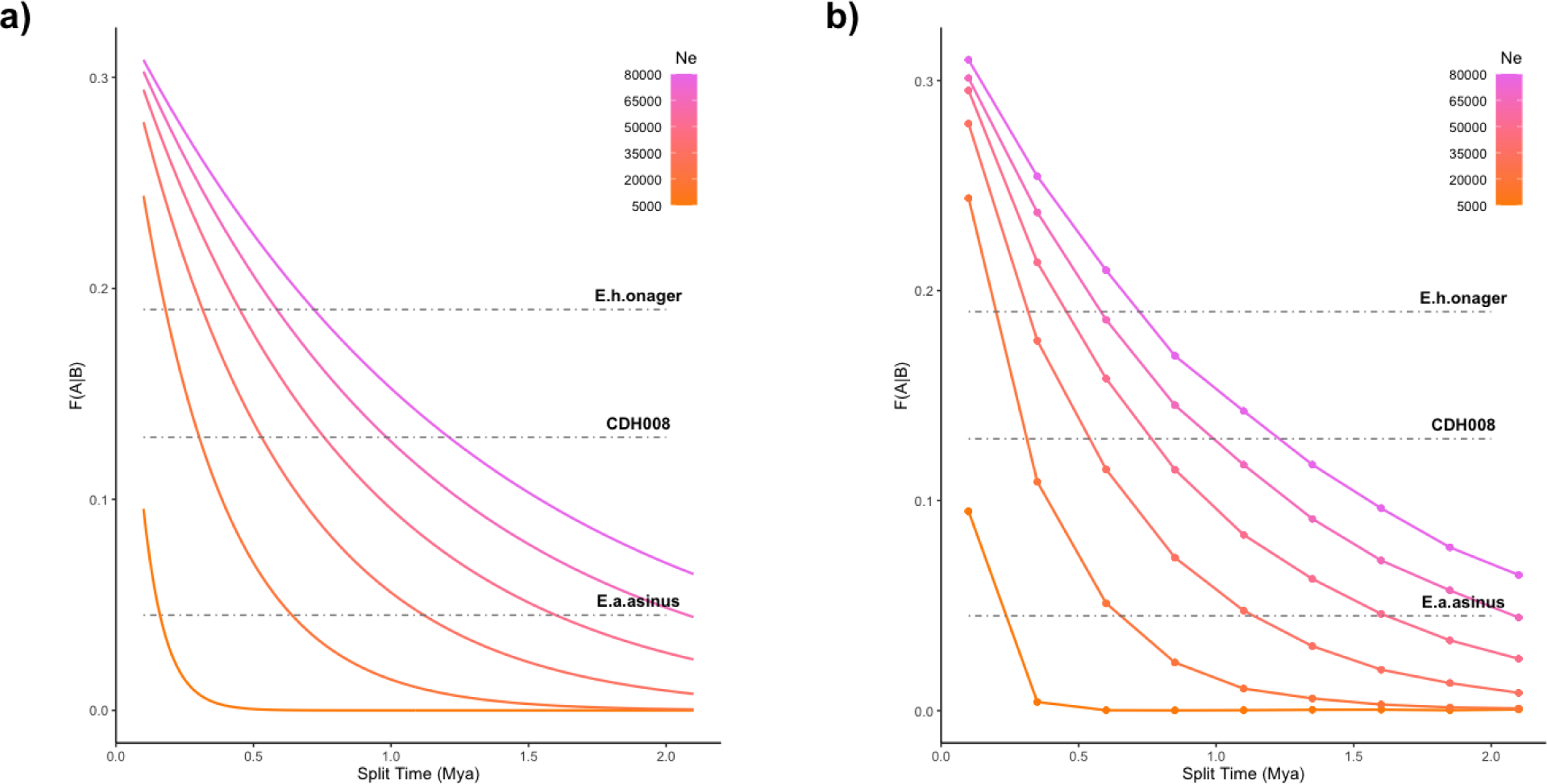
Population split time estimation of Anatolian wild ass (CDH008), *E. a. asinus* and *E. h. onager* from Asian wild ass based on the F(A|B) statistic. F(A|B) is the probability of observing a derived allele in the Anatolian wild ass, *E. a. asinus* or *E. h. onager* at the positions where the Asian wild ass genome is heterozygous (Methods). Observed and expected F(A|B) values, calculated using a) e^−T/(2N)/3^ and b) population genetic simulations, are given for divergence times ranging from 100kya to 2 mya, and for population sizes ranging from 5k to 80k.

### Severely depleted genetic diversity in the Anatolian wild ass signals population decline

To our knowledge, these three Anatolian individuals are the latest directly dated hydruntines known, if not the latest recorded hydruntines to date (Guimaraes et a., 2020; Crees & Turvey, 2014). Zooarchaeological evidence suggests that by the 3^rd^ millennium BCE the European population had already gone extinct, while the population was possibly dwindling or lost in Southwest Asia, including in Anatolia, Iran, and the Caucasus (Boulbes and van Asperen, 2019; Crees and Turvey, 2014). We thus asked whether we might detect signatures of a stark population decline in our genetic data.

We first estimated runs of homozygosity (ROH), thus measuring past inbreeding that can be caused by shrinking population size (Ceballos et al., 2018). We called ROH in the highest coverage Anatolian wild ass genome (cdh008) and compared these ROH estimates with those from genomes of other asses, after down-sampling all to the same coverage (Methods). We observed higher numbers of ROH in the African asses (*E.a.asinus* and *E.a.somalicus*) (Figure 6A), as expected, since these populations are known to have experienced recent bottlenecks (Jónsson et al., 2014; Renaud et al., 2018). On the other hand, *kiangs, kulans* and *onagers* showed relatively low ROH loads, implying higher diversity than their African counterparts. In turn, cdh008 carried a high number of short runs with complementing long runs, highly similar to the African genomes.

**Figure 6.**
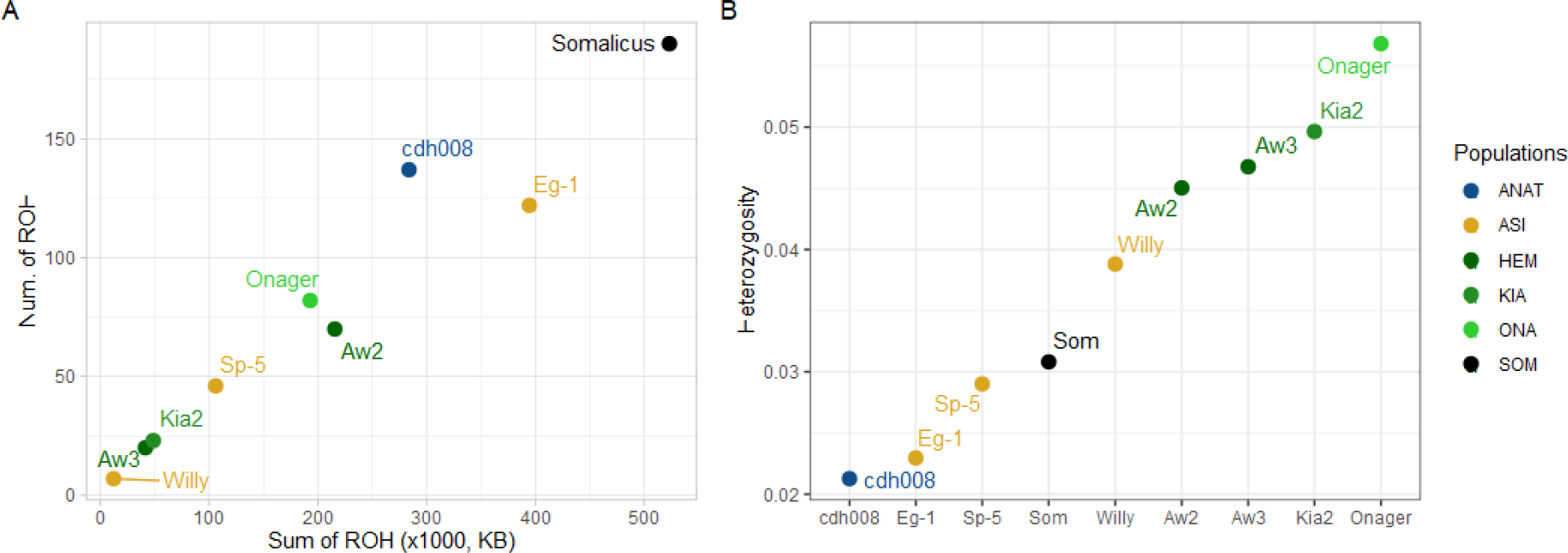
A) Total number versus the total length (sum) of runs of homozygosity (ROH) tracks over 1.5 Mb. B) Heterozygosity calculated across 2,146,416 transversion sites.

We investigated this further by estimating heterozygosity from *de novo*-called transversion variants across the same genomes. Hemiones had the highest heterozygosity, followed by donkeys and the Somali wild ass. The cdh008 genome harbored the lowest heterozygosity, even below the African group known to have undergone several bottlenecks (Figure 6B). Combined with ROH analysis, these observations suggest that the Anatolian population was in strong decline already by the 1^st^ millennium BCE, which is also supported by its rare occurrence in the archaeological record.

### Scans for possible hydruntine-specific positive selection and relaxation in coding regions

We investigated possible hydruntine-specific protein-coding adaptive changes by comparing the cdh008 genome against those of Asian and African ass lineages. We used two methods. First, we ran *PAML codeml* (Yang, 2007) on 15,985 open reading frame alignments, using the tree in Figure 2B (Methods). In total, 179 genes showed only nominally significant evidence for positive selection along the cdh008 branch [chi-square test nominal p<0.05, calculated following Zhang et al., 2005] (Supplementary Table 7), although none were significant after multiple testing correction. We also identified similar numbers of putatively positively selected genes on the Asian branch (190 genes). Gene Ontology (GO) analysis did not yield any significantly enriched GO categories among these genes, neither for the Anatolian branch nor the Asian branch (Fisher’s Exact Test p-value>0.05 after Benjamini-Hochberg false discovery rate correction).

Because *codeml* analysis is expected to have low power when lineages are closely related, we attempted to bolster the scan by using a 3-lineage test and including polymorphism data. Namely, we searched for coding sequences where the Anatolian wild ass was differentiated from both Africa and Asia, possibly reflecting positive selection, while Africa and Asia had not diverged, presumably due to purifying selection in the latter lineages. For this, we combined the neutrality index based on the McDonald-Kreitman table, i.e. (*D^N^*/*D^S^*)/(*P^N^*/*P^S^*) (Stoletzki & Eyre-Walker, 2011) with a framework inspired by the population-branch statistic (PBS) (Yi et al., 2010) (Methods). We used polymorphism data (*P^N^* and *P^S^*) only from African and Asian asses, as the Anatolian sample was too small to obtain reliable polymorphism calls (we thus use polymorphism here only for estimating purifying selection). We call this statistic the Pairwise McDonald-Kreitman (PMK).

Using PMK we scanned the equid genome for outlier genes where Anatolian asses had diverged from the other two groups above the genome average. Out of 17,327 genes analyzed (Supplementary Table 8), we identified 227 with a *Z*-score > 3, and 114 of these passed the Benjamini-Hochberg multiple testing correction at p < 0.01, with higher *D^N^* on the Anatolian lineage than the other ass lineages (Supplementary Table 8; Supplementary Figure 13). We again did not identify enrichment in GO categories after multiple testing correction (p>0.05), but noted that the *PAML codeml* and PMK outlier lists shared 7 genes after PMK multiple testing correction (1.3 expected, Fisher’s exact test p=3.1×10^−4^): *CD86, GPRC5A, KMO, LOC100064229, LOC102148548, MACF1, ODF4, ZNF45* (Supplementary Table 9).

Finally, we asked whether small effective population size, as indicated by the high ROH load, may have led to elevated non-synonymous mutation load in the hydruntine lineage due to relaxed selection. Comparing *PAML*-estimated *dN*/*dS* ratios for cdh008, Asian, and African genomes (Methods), we found slightly but significantly lower values in cdh008 than the other two genomes (Supplementary Figure 14).

## Discussion

We have presented analyses of three ancient genomes of Anatolian wild asses radiocarbon dated to the 1^st^ millennium BCE. All three carried a putative hydruntine-specific mitochondrial haplotype (*H1*; Bennett et al., 2017), and phylogenetic analyses of their mtDNA sequences consistently clustered these individuals together with morphologically-identified hydruntine (*E.hydruntinus* or *E. hemionus hydruntinus)* sequences with full support. The three genome profiles also fell clearly outside the diversity of known African or Asian ass genomes, while being similar within themselves. Together, the evidence strongly indicates that the three Anatolian wild ass individuals belonged to the extinct hydruntine *E.hydruntinus/E.h.hydruntinus*.

Although it has been termed the “European wild ass”, the hydruntine range included Southwest Asia and possibly even North Africa (reviewed in Boulbes and van Asperen, 2019). In Anatolia, hydruntines were not uncommon in zooarchaeological assemblages until the Bronze Age (Arbuckle, 2013; Arbuckle & Öztan, 2018; Bennett et al., 2017), although their presence largely depends on the site and past subsistence strategies (Martin & Russell, 2013). At Neolithic Çatalhöyük, for example, small and medium-sized equids (*E. hydruntinus/hemionus*) are evidenced by bones in daily and social contexts (Pawłowska, 2020) and in art, although being relatively rare. In Çatalhöyük art, they appear in one set of wall paintings, while phalanges were used as raw material for making idols, one of which is known as the first bone figurine from the site (Martin and Russell, 2013; Pawłowska and Barański, 2020).

Until now, the last hydruntine detected on a genetic basis in Anatolia was dated to ∼2,200 BCE (Guimaraes et al., 2020). In Iran, osteologically-identified specimens were dated to the 2^nd^ millennium BCE (Mashkour et al., 1999). Our three Anatolian wild asses, dated to early/middle 1^st^ millennium BCE, are thus the last hydruntines yet identified.

Our study further reveals a number of novel insights into the phylogeny of hydruntines, represented by the three Anatolian genomes. First, using genomic evidence we confirm that Asiatic hemiones and hydruntines were sister taxa to the exclusion of African asses, as suggested earlier based on osteological data (Burke et al., 2003; Eisenmann & Mashkour, 1999; Orlando et al., 2006) and mtDNA analyses (Bennett et al., 2017; Catalano et al., 2020; Orlando et al., 2006, 2009). Thus, we may speak of Late Pleistocene and Holocene Eurasiatic wild asses as a broadly coherent evolutionary group.

We do not find support, either using mtDNA or nuclear genome analyses, that hydruntines were a sub-branch within the known Asiatic hemione diversity; instead, they appear to be the earliest diverging branch of Eurasian wild asses (Figure 1B). Asiatic wild asses, represented by the hemione, kiang, onager and hemippe genomes used here, appear monophyletic in the full mtDNA trees, nuclear genome trees, and in D-statistics, to the exclusion of hydruntines. This result, along with the apparently distinct phylogenetic histories of hydruntines and hemiones (Figure 4), as well as potential differences in their major phenotypes (e.g. the hydruntine short muzzle) together support the idea that hydruntines, instead of being considered as a hemione sub-branch (Orlando et al., 2009; Bennett et al., 2017), may deserve an independent taxonomic status from other hemiones (Boulbes and van Asperen, 2019; van Asperen, 2012). However, because our genome dataset did not include the full diversity of hemiones (e.g. the Indian khur and Central Asiatic wild ass genomes are not yet sequenced), because we lacked the information about full hydruntine diversity through ages, and also because our Anatolian samples represented a population that appears to have undergone a severe bottleneck, reciprocal monophyly between hemiones and hydruntines may still be considered an open question. It was proposed previously that the European hydruntine may have split from the Central Asiatic wild ass populations and evolved independently in Europe before colonizing Anatolia in the Late Pleistocene/Early Holocene (Bennett et al, 2017). Genomes of European hydruntines, as well as recent or extant asses from Central and South Asia could help fully comprehend the evolution of this taxa. Overall, our hydruntine genomic data strongly support a long-time independent evolution of the hydruntine and the other south-west Asiatic wild asses, compatible with previous interpretations (Bennett et al. 2017).

Our finding of genes with elevated rates of non-synonymous changes in the hydruntine lineage is also consistent with the idea that the hydruntine lineage carried distinct adaptations (although this was not tested against a null model of coding-sequence evolution). A notable candidate here was *MACF1*, a highly pleiotropic gene encoding for a microtubule-interacting and cell migration-related protein. Missense variants in *MACF1* have been associated with diverse phenotypes, including brain development defects (Dobyns et al., 2018) and Parkinson’s disease risk (Wang et al., 2017) in humans, and carcass weight in cattle (Niu et al., 2021). The locus has also been reported to show signatures of Neanderthal-to-human adaptive introgression (Gower et al., 2021), as well as signatures of recent positive selection in the buffalo (Sun et al., 2020) and cattle (Maiorano et al., 2018). Given its signature of non-synonymous divergence on the hydruntine lineage, we speculate that substitutions in *MACF1* might explain some of the peculiar features of the hydruntine.

Our results also indicate gene flow from hydruntines into Middle Eastern hemiones, i.e. the onager of Iran and the hemippe of Syria. This can be inferred from the asymmetric relationships between hydruntine and Middle Eastern vs. Gobi/Tibet hemiones, with hydruntines being on average closer to the former (Supplementary Figures 7-9; Supplementary Tables 5-6). Likewise, the hydruntine is closer to the modern onager and 19^th^ century hemippe genomes than to Iron Age onagers and a hemippe from the Neolithic period, respectively (although the result is variable between the SNP panels) (Supplementary Figures 10-11; Supplementary Tables 5-6). These observations also imply that gene flow occurred from hydruntines into Middle Eastern hemiones in recent times. This is because, assuming no strong population structure within onagers or hemippes, gene flow from the onager to hydruntines would not lead to asymmetric relationships with different onagers or hemippes. Such gene flow between hydruntines and hemiones may not be surprising, given evidence for the widespread introgression among equids (Jónsson et al., 2014), and the possibility that hydruntines may have lived in sympatry with these Middle Eastern hemiones in Southwest Asia during the Holocene (Crees and Turvey, 2014; Eisenmann & Mashkour, 1999; Orlando et al., 2006). Our data thus suggest the lack of strict reproductive isolation between hydruntines and Middle Eastern hemiones. This could be considered a reason for grouping hydruntines under *E. hemionus*, even though the limited extent of these admixtures suggests that these wild asses were distinct enough to not fully interbreed, supporting the idea that the hydruntine had already experienced a large portion of the speciation process separating them from Middle Eastern wild asses.

Finally, our results shed light on the extinction dynamics of the hydruntine in Southwest Asia. In a recent study the last genetically detected Anatolian hydruntines dated to the end of the 3^rd^ millennium BCE (Guimaraes et al., 2020). Although we find that hydruntines survived at least two more millennia, our ROH and heterozygosity analyses on the 1^st^ millennium BCE genome from Çadır Höyük do suggest a dramatic reduction in their effective population size in that period on a par with the critically endangered African wild ass (Moehlman & Kebede, 2014), which would have dramatically increased the risk of extinction (Newman and Pilson,1997). The decline itself could have various non-exclusive causes, including habitat fragmentation as a result of human activity and/or climatic events, or human predation (Bennett et al., 2017; Boulbes & van Asperen, 2019; Cai et al., 2021; Crees and Turvey, 2014; Spassov & Iliev, 2002). It is notable here that the remains of the three individuals studied here were recovered from residential or midden contexts (Archeological Supplement Table 2) and were likely consumed by humans. Meanwhile, the lack of an elevated *dN*/*dS* signal in the cdh008 genome appears surprising, given high levels of damaging variant loads in natural populations subject to strong bottlenecks (e.g. Robinson et al. 2021) or in domesticates (reviewed in Bosse et al. 2018). One possibility is that the reduction in the hydruntine population size may have been too recent to visibly impact functional variation.

Another question remaining is how much longer hydruntine populations survived in Southwest Asia, beyond the individuals we sampled from Anatolia. The trend of higher affinity of hydruntines to the modern-day than to Iron Age onagers implies that hydruntines may have admixed with onagers after the Iron Age (Supplementary Figure 12). Interestingly, classical authors such as the Roman historian Strabo (*Geography,* Book XII), Varro (*De Re Rustica*, Book II), and Pliny the Elder (*Natural History*) mention the presence of “wild asses” (*onagri*) in the vicinity of ancient Late Tatta (modern-day Tuz Gölü in Central Anatolia), and also in the Anatolian dry grasslands of Lycaonia, Garsauira, and Bagadania. Therefore it seems possible that these may be referencing hydruntine populations surviving on the central Anatolian plateau (in the Konya-Ereğli plain, Cappadocia, and/or Kayseri) into the first millennium AD. Our results underscore the need for further archaeogenomics-based taxonomic assignment of non-caballine equid material from Southwest Asia. This would help fully fathom the extinction dynamics of the hydruntine.

## Material and Methods

### Archaeological samples

We studied skeletal samples identified as equid, 11 from Çatalhöyük and four from Çadır Höyük. Çatalhöyük is a major Ceramic Neolithic period site in Central Anatolia, but its upper layers have also yielded remains dating to Bronze and Iron Ages, and later periods [Hordecki, 2020; Pawłowska, in press]. Çadır Höyük has demonstrated continuous occupation from the Middle Chalcolithic to the Byzantine Era (early fifth millennium BCE to fourteenth century CE) (Ross et al. 2019; Steadman et al. 2019a, 2019b). See the Archaeological Supplement for further information on the sites and the archaeological material.

### Radiocarbon dating

We radiocarbon dated all three equid samples which showed wild ass genetic signatures. For each sample, approximately 3 grams of bone or tooth material was cut using non-carbon based discs attached to a Dremel tool and sent to the TÜBİTAK-MAM (Gebze, Turkey) AMS facility for carbon-14 dating. The dates were calibrated using the R package “*Bchron*” using the IntCal20 (Reimer et al., 2020) curve.

### Ancient DNA isolation and sequencing

All pre-PCR experiments were conducted in the METU Ancient DNA Clean Room, in a dedicated laboratory for aDNA research, located in a different building from the post-PCR laboratory. Samples were subjected to a standard ancient DNA isolation protocol (Dabney et al., 2013), with minor modifications. In brief, surfaces of archaeological samples were cleaned with damp paper cloth, 100-200 mg of bone or tooth sample was cut using discs attached to a Dremel, were pulverized with mortar and pestle, and transferred into 2 mL screw-top tubes. Each sample powder was treated with 1 mL extraction buffer (0.45 M EDTA and 0.25 mg/mL Proteinase K) in a 37° C rotating incubator for 18 hours. Tubes were centrifuged and supernatants were added to reservoirs containing 13 mL of binding buffer (5 M Guanidine Hydrochloride, 40% (vol/vol) Isopropanol, 0.05% Tween-20, 90 mM Sodium Acetate) to bind DNA fragments to Qiagen spin columns. DNA fragments were washed twice with a washing buffer and fragments were collected after two consecutive elution steps into 50 µL Qiagen EB buffer. Blunt-end ancient DNA libraries were prepared using Meyer and Kircher protocol (Meyer & Kircher, 2010) with a single indexing approach. All experiments on Çadır Höyük and Çatalhöyük samples were conducted on different days with fresh reagents prepared for each archaeological site.

Libraries were sequenced on the Illumina HiSeq 4000 platform. Using 47 thousand - 79 million reads (median 20 million) obtained in initial low-coverage sequencing, we identified three equid samples, two from ÇadırHöyük (cdh008, phalanx; cdh010, calcaneus) and one from Çatalhöyük (chh003, tooth), that displayed a wild ass-related genetic signature. We further sequenced cdh008 three times, cdh010 and chh003 one more time.

### Data preprocessing

The data was processed following (Kılınç et al., 2016) and (Günther et al., 2015). Raw sequencing reads were demultiplexed, adapter sequences were removed and paired-end sequencing reads were collapsed using *AdapterRemoval* (v. 2.3.1), requiring a minimum of 11 bp overlap between pairs (Schubert et al., 2016). Collapsed *.fastq* files were aligned to horse reference genome (EquCab2.0; including 31 autosomal chromosomes, the X chromosome, and the mitochondrial genome) (Wade et al., 2009) using the *aln* module of *BWA* software (v. 0.7.15) (Li & Durbin, 2009) with the parameters *“-n 0.01 -o 2”* and disabled the seed with *“-l 16500”*. All alignment files of the same individual were merged using *SAMtools* (v. 1.9) *merge* (Li et al., 2009), PCR duplicated reads were removed using *FilterUniqueSAMCons.py* (Meyer&Kircher, 2010) and reads with length <35 bp were discarded. Reads with a mismatch to fragment length ratio >10% were also removed.

### Authentication of genetic data and trimming

We verified the authenticity of the genetic data using *PMDtools* (v. 0.60) (Skoglund et al., 2014), which quantifies post-mortem damage (PMD) signals at both ends of mapped reads, characteristic of ancient DNA (Briggs et al., 2007). PMD profiles for each sample were generated using the *PMDtools “--deamination”* parameter. Deamination signals of cytosine to thymine transitions at 5’-most positions were 0.31, 0.29 and 0.36 for cdh008, cdh010 and chh003, respectively. Average read lengths were 74 bp for cdh008, 75 bp for cdh010 and 67 bp for chh003, again supporting authenticity (Pedersen et al., 2014).

Reads obtained from libraries were trimmed 10 bp from both ends using “trimBAM” command of the *bamUtil* software (Jun et al., 2015) to remove residual deamination. We noted that PMD also presents a confounding effect on genotyping at transversion positions, for example by lowering the probability of observing C at C/A polymorphisms.

### Complete mitogenome sequence

We generated a complete mitogenome sequence of our highest coverage sample cdh008 (coverage: 140.04X) by consensus calling of the most common bases using Geneious Prime 2022.2 with default parameters. Most of the unidentified bases (Ns: 2% of the whole sequence) were observed near the end of the sequence. The sequence was deposited to the NCBI Genbank with accession no: OP448588.

### Taxonomy and sex determination

We first investigated the taxonomic identity of these three samples using the *Zonkey* pipeline implemented in *PaleoMix* (v. 1.2.14) (Schubert et al., 2017). *Zonkey* generates a PCA (Patterson et al., 2006), *TreeMix* (Pickrell& Pritchard, 2012) and mitochondrial phylogeny graphs based on a reference panel consisting horse, zebra, donkey, and Asian wild ass (hemione) genomes. SNP counts used in *Zonkey* analysis were 163,921 for cdh008, 158,448 for cdh010 and 162,739 for chh003. All three samples were clustered together with the *Equus hemiones onager* and *Equus kiang* (Asian wild ass) group, confirming their Eurasian wild ass status. Sex estimation was also performed using *Zonkey*, based on autosomal versus X chromosome coverages. For chh003, X chromosome coverage was approximately half the autosomal coverage, suggesting this sample was a male individual, while the data for cdh008 and cdh010 showed similar coverage in autosomes and the X chromosome, suggesting they were female.

### Phylogenetic analyses of mtDNA data

Phylogenetic analysis was carried out using mtDNA consensus sequences generated from *BAM* files using *angsd* (v. 0.934)(Korneliussen et al., 2014) with *“-doFasta 2”*, *“-minQ 30”*, *“-minMapQ 30”* and *“-setMinDepth 3”* parameters.

*MEGA X* (Kumar et al., 2018) was used to construct phylogenetic trees using maximum likelihood and neighbor-joining. To this end, a 361 bp fragment of the D-loop in the control region of the mitochondrial DNA, as well as a 13,042 bp long consensus sequence of the whole mitochondria were used (Genbank). For the maximum likelihood analysis using 138 D-loop sequences (Supplementary Table 6), the substitution model was chosen as Hasegawa-Kishino-Yano (HKY) with a Gamma-distributed rate (no invariant sites) using *jModeltest* (v.0.1.1) (Guindon & Gascuel, 2003; Posada, 2008). A maximum composite likelihood model was used for the neighbor-joining tree construction. The nodal support was evaluated by 1000 bootstrap replicates for both methods. For the median joining (MJ) tree, 121 partial D-loop samples were trimmed to 249 bp based on the shortest fragment in the dataset. The MJ network was constructed with *NETWORK* v.5 maximum parsimony post processing (http://www.fluxus-engineering.com).

The mitochondrial DNA divergence times of chosen equid species were estimated with the full mitochondrial DNA sequence using Bayesian inference with *BEAST* (v. 1.8.4) (Drummond et al., 2012). For this analysis, the best substitution model was found to be GTR with a Gamma distributed rate and invariant sites using *jModeltest* (v. 0.1.1). An uncorrelated relaxed clock with a lognormal distribution and Yule speciation was assumed. The convergences were visualized using *Tracer* (v. 1.6) (Rambaut, 2014). The best tree was found and drawn using *TreeAnnotator* (v. 1.8.4) (Drummond et al., 2012) and *FigTree* (v. 1.4.3) (Rambaut, 2014). *Equus caballus* was used as an outgroup and the molecular clock was calibrated using a normal prior of 4.25±0.125 Mya for the divergence of *E. caballus* (Vilstrup et al., 2013). The analysis was run for 600,000,000 steps, logging every 60,000 states and discarding the first 10% states as burn-in, resulting in effective sample sizes (ESS) greater than 1000 for all traces. The mitochondrial divergence between African and non-African asses was estimated as 3.57 [1.18-4.23] mya, which is higher than but includes previous estimates (*e.g*. 2.62 mya) (Vilstrup et al., 2013).

### SNP panel construction

A genomic variation dataset was constructed using 12 modern-day high coverage (>5x) genomes (Huang et al., 2015; Jónsson et al., 2014; Renaud et al., 2018; Wang et al., 2020), of which six were domesticated donkeys (*Equus asinus asinus*) and six were Asian wild asses (three *Equus hemiones hemionus*, one *Equus hemiones onager*, two *Equus hemionus kiang*)(Supplementary Table 3). The *E.h.onager* sample was traced back in their lineage to confirm its species. It revealed that this individual’s (studbook#818) father (studbook #682) was an onager with known origin while mother was a non-specified onager. Their father’s (studbook #682) father was a non-specified onager and mother was a zoo onager (studbook #618). The lineage of #618 individual was captured from Iran in the wild between 1955-1973; 6-7 generations before the birth of #682. Two horse samples were included as an outgroup (Der Sarkissian et al., 2015; Orlando et al., 2013) (see Supplementary Table 3). First, we performed *de novo* SNP calling using *angsd* (v. 0.934)(Korneliussen et al., 2014) from donkey and wild asses separately. We limited *de novo* SNP calls to positions with a minimum depth of 4, and required that the SNPs were non-missing across all the individuals. A total of 31,592,013 and 38,029,882 SNPs were thus called from donkeys and wild asses, respectively. Limiting these to autosomal positions, applying a minor allele frequency filter of MAF > 0.1, a Hardy-Weinberg filter of “1e-30” and keeping only biallelic transversion positions yielded 2,146,416 SNPs, which we refer to as the main SNP panel.

To confirm results from this main SNP panel using a published alternative, we used a second SNP panel of equid variation as selected in Bennett et al (2022). This panel was derived from the panel used in the *Zonkey* algorithm that was established from a single genome of each of 9 Equids species or subspecies, including horses, African and Asiatic wild asses and zebras (Schubert et al. 2017). This set of SNPs had been previously filtered to eliminate all SNPs that fell within repeats in the reference genome based on the assumption that repeats are more prone to artifactual mapping with ancient DNA fragments, in particular when mapping to the genome of a distantly related species (Bennett et al 2022). We further filtered this set to keep positions polymorphic in asinus and hemionus and used only transversions, leaving us with 2,408,064 SNPs. We refer to this as the secondary SNP panel.

### Genotyping and dataset construction

We genotyped both modern-day and ancient genomes using either of the SNP panels. All modern-day individuals were genotyped using *SAMtools (v. 1.9) mpileup* module in diploid fashion. Ancient individuals were genotyped using *sequenceTools (*v. 1.4.0.5) (https://github.com/stschiff/sequenceTools) using the option “-randomHaploid”. The *pileupCaller* module of this program was run on ancient samples, genotyping only the positions in the SNP panel. Ancient genomes were pseudohaploidized by randomly choosing a single read and using its allele as the individual’s genotype. Genotypes of modern-day and ancient individuals were merged using *PLINK* (v. 1.90) (Purcell et al., 2007) and *EIGENSOFT* (v. 7.2.1)(Patterson et al., 2006) to form the final versions of the dataset. We thus prepared two datasets: one we refer to as the main dataset, the other as the secondary dataset. These were used in all population genetic analyses except for selection scans and the CDS-based phylogenetic tree construction.

### *f*^3^-statistics

We performed outgroup-*f*^3^ statistics on the autosomal chromosomes dataset using *popstats* (Skoglund et al., 2015) with *“--not23”* and *“--f3 vanilla”* parameters, and using the *PLINK* (v. 1.90) *.tped/.tfam* file format. Analyses were run in the forms of *f^3^(Horse;Anatolia,Modern)*, *f^3^(Horse;Modern,Modern)*, *f^3^(Horse;Anatolia,Anatolia)* and *f^3^(Horse;Anatolia,AncientHemione)* in a pairwise manner, where *Horse* represents two horse genomes used as outgroup, *Anatolia* represents one of three Anatolian ancient wild ass individuals reported in this article, *Modern* represents a modern-day donkey, Somali wild ass, or Asian wild ass genome, and *AncientHemione* represents the three published ancient hemione and three ancient hemippe genomes mentioned earlier (Supplementary Table 3). A heatmap graph was generated from the *f*^3^ results using the *heatmap.2* command of the R package *gplots* (https://cran.r-project.org/web/packages/gplots/index.html).

### Multidimensional scaling

A genetic distance matrix was generated by subtracting pairwise outgroup-*f*^3^ values from 1. This matrix was used in multidimensional scaling analysis by running the *“cmdscale”* command in R with parameters *“eig=True”* and *“k=2”*. The first two dimensions were visualized in R.

### D-statistics

Gene flow and genetic affinity between ancient and modern groups were inspected with D-statistics using main and secondary dataset in *EIGENSTRAT* file format (*.geno/.ind/.snp*), running the *“qpDstat”* module of *AdmixTools* (v. 7.0) (Patterson et al., 2012) with default parameters. The analysis was run on all possible trio combinations, using horse as an outgroup. We limited the analyses to pairs with sufficient numbers of overlapping SNPs (for example excluding historical hemippe individual, Hm_1892). Significance of each comparison was inspected using the *Z*-score (the ratio of the D-statistic to the standard error, the latter estimated using a jackknife procedure). Results with absolute *Z*-scores > 3 (corresponding to roughly p<0.001) were defined as nominally significant. Results were visualized using the R package, gg*plot2* (Wickham, 2016).

### Autosomal CDS tree

Following (Chen et al., 2021), we created a phylogenetic tree of equid lineages based on autosomal protein coding sequences (CDS); the choice of using CDS is to facilitate alignment and data processing. CDS of three *E.h.hemionus*, two *E.h.kiang*, one *E.h.onager*, three *E.h.hemippus,* three *E.hydruntinus* reported in this study, one *E.a.somalicus*, two *E.a.asinus* and one *E.caballus* (as an outgroup) (Supplementary Table 3) were extracted from whole genome data, based on the annotation file from NCBI (In case of overlapping genes, the longer one was taken into account). Selected CDS genes were extracted from bam files by *angsd* (v. 0.934) (Korneliussen et al., 2014) with the base with the highest effective depth (“-doFasta 3” option), and requiring a minimum base quality of 30 and also a mapping quality of 30 (“-minQ 30” and “-minMapQ 30” options, respectively). Resulting fasta files were aligned and trimmed to the same size for each chromosome. A final CDS dataset was constructed by merging all chromosomes and filtering all transitions. CDS selection, aligning-trimming, merging and filtering the transitions were all performed by custom python scripts (filter_gene.py, trim_py merge_py, and remove_transitions.py respectively). A maximum likelihood tree was generated with this CDS dataset using *RAxML*v. 8.2.12 (Stamatakis, 2014) with the following strategy: A preliminary tree was constructed using the GTRCAT model, 200 bootstrapping replicates were applied on the preliminary tree using the GTRGMMA model, and a final tree was generated by overlaying the calculated bootstraps on the preliminary tree.The final tree was visualized by R package ape v. 5.6.2 (Paradis and Schliep, 2019) and R package phytools v. 1.0.3 (Revell, 2012).

### Divergence time estimation

In order to estimate the divergence time between the hydruntines and asian wild assess, we used the F(A|B) statistic (Green et al. 2010), which calculates the probability of observing a derived allele in individual A, at the sites where individual B is heterozygous. For our case, individual A corresponds to our best quality hydruntine cdh008, and individual B corresponds to the kiang sample Kia2 (Wang et al. 2020), representing the Asian lineage. In total n=2,626,614 heterozygous positions were determined in the Kia2 genome. The alleles were sorted as ancestral or derived according to the horse genome (EquCab 2.0). Genotype calling in cdh008 was carried out using *samtools* (v. 1.9) *mpileup* (Li et al., 2009), with a minimum read and base quality of 30. The probability was calculated by choosing a random allele at each site. This probability was then computed for different divergence times and population sizes using the formula e^−T/(2N)^/3 (Mualim et el., 2020) and simulations using *msprime* (v. 1.0)(Baumdicker et al., 2022), with a generation time of 8 years, a mutation rate of 7.242×10^−9^ and a recombination rate of 1cM/1Mbp (Orlando et al. 2014). We simulated 100 chunks of 1Mb, and calculated the mean F(A|B). Using the formula *H*=4*N^e^µ*, we estimated *N^e^* for Kia2 as 62,319.3. Expected F(A|B) values were calculated for divergence times ranging from 100kya and 2.1mya, and for population sizes ranging from 5k to 80k. We determined the expected F(A|B) values which span the observed F(A|B) value to find the estimated time of population split. For comparison, we also estimated split times of *E. h. onager* and *E. a. asinus* each from *E. kiang*, again using F(A|B). We also repeated the F(A|B) calculation using only transversions (n=811,814) instead of all SNPs, which resulted in a highly similar estimate (0.145 using all SNPs vs. 0.149 using transversions only). We note that using transversions limits the chance of homoplasy and misidentification of the ancestral state.

### Runs of homozygosity

Runs of homozygosity (ROH) of representative samples for each group of *E.a.asinus*, *E.h.kiang*, *E.h.onager*, *E.h.hemionus*, *E.a.somalicus* and “cdh008” for the Anatolian wild ass, (or *E.hydruntinus*) (Supplementary Table 3) were calculated from equal coverage *BAM* files. For this, we downsampled all *BAM* files to 6.38x to match the lowest coverage case, cdh008. To calculate ROH, we used *PLINK* (v. 1.90)(Purcell et al., 2007) with the following parameters: “*--chr-set 34 --homozyg -homozyg-snp 30 - homozyg-kb 300 -homozyg-density 30 -homozyg-window-snp 30 -homozyg-gap 1000 -homozyg-window-het 3 -homozyg-window-missing 5 -homozyg-window-threshold 0.05*”. The resulting data were merged into a single file and visualized using R. Our recent work has shown that ROH can be called efficiently using this approach from >5x genomes (Ceballos et al. 2021).

### Genome-wide heterozygosity estimation

In order to estimate genome-wide heterozygosity for modern-day and ancient equid genomes, we first downsampled *BAM* files from modern-day equids, namely three *E.a.asinus*, one *E.a.somalicus*, one *E.h.onager*, two *E.h.hemionus* and one *E.h.kiang* to 6.38x coverage, i.e. the coverage of the Anatolian cdh008. Using these downsampled *BAM* files and that of cdh008, we called de novo variants in each genome at all positions with 5-13x depth using *samtools* (v. 1.9)(Li et al., 2009). We then determined heterozygous loci for each genome, and further filtered these sites to only keep transversion positions to avoid PMD-related false heterozygosity calls. Finally, the total number of heterozygous sites were divided to the number of all positions with 5-13x depth in the downsampled genomes. Results were visualized using the R package, gg*plot2* (Wickham, 2016).

### Constructing gene sets for selection scans

Here our goal was first to create protein coding gene sequences for each of the Asian, African and Anatolian wild ass lineages, where each sequence carries fixed nucleotide changes (substitutions) in that lineage. For this, we generated a VCF file including Asian, African and Anatolian ass genomes aligned to the EquCab2 reference genome and called *de novo* SNPs only for these individuals using *angsd* (v. 0.934)(Korneliussen et al., 2014). The Anatolian group included the cdh008 genome, the Asian group included the genomes Aw1, Aw2, Aw3, Kia1, Kia2 and Onager, while the African group included the genomes Sp-5, Eg-1, Ch-by1, Ke-14, Ky-5, Ir-5 and Somalicus (Huang et al., 2015; Jónsson et al., 2014; Wang et al., 2020). For each of the three groups, we chose biallelic SNPs genotyped as alternative allele at least once, but never as the reference allele, using *bcftools* (v. 1.11) *view* (Danecek et al., 2021) with options ‘-c 1 -e ‘GT==“RA”|GT==“RR””. These represent possible fixed substitutions on each lineage (with the caveat that the single Anatolian genome used is represented by a relatively low coverage genome, and thus its putative substitution set may include a certain frequency of polymorphic sites). For each of the three groups, the selected substitutions were inserted into the EquCab2 reference *FASTA* file using the *vcftools* (v. 0.1.16) (Danecek et al., 2011) *vcf-consensus* programme, yielding three separate *FASTA* files. From each of the resulting *FASTA* files, we extracted protein coding regions with the *cufflinks* (2.2.1) *gffread* (v. 0.12.7)program (Pertea&Pertea, 2020) with the “-x” option, using gene annotations from the NCBI Annotation Release 102. We call this the Gene Set 1.

We created another set, Gene Set 2, representing polymorphic sites within lineages. We used a similar approach, but this time we selected the positions where we observed at least one reference and at least one alternative allele in the population. This was performed on Asian wild ass and African ass groups, separately, but not on the Anatolian genome cdh008 due to lack of coverage.

### PAML codeml analysis

To run *PAML* (v. 4.10.6) *codeml*, we created multiple alignment files for each coding sequence (N=32,988) using the three representative sequences described above (Gene Set 1), as well as the horse reference to be used as an outgroup, using an in-house script.

Among these multiple alignment files including alternative forms of a gene, 58 were excluded because the sequence lengths were not a multiple of three, 304 were excluded because they lacked stop codons, 476 because they contained premature stop codons in all four lineages, while 909 were excluded because they lacked stop codons or contained premature stop codons or were not triplets in some but not all lineages. The remaining 31,241 multiple alignment files were further filtered to only contain the longest transcript of a gene if the gene had multiple isoforms. When there were multiple alternative isoforms of the longest transcript of the gene, the first one in alphabetic order was selected. All in all, the remaining 15,985 multiple alignment files were subjected to the branch-site test for positive selection with the *PAML codeml* program, using parameters “model=2” and “nssites=2” (Zhang et al., 2005). In the null model ⍵ is fixed whereas in the selection model it is allowed to vary both among sites and branches. The likelihood values of the two models were compared according to the branch-site test of positive selection (Zhang et al.,2005). Gene Ontology Analysis testing enrichment of a putatively selected set of genes relative to all tested genes was conducted with the R *topGO* package (Alexa & Rahnenführer, 2020) [R package version 2.38.1.]

### Pairwise McDonald-Kreitman analysis

For calculating the Pairwise McDonald-Kreitman (PMK) statistic, we used the Gene Set 2 described above. We again considered the longest transcript per gene as described above and removed CDS that were not a multiple of three in length. For each gene, we first counted the numbers of synonymous substitutions (*D^S^*) and nonsynonymous substitutions (*D^N^*) between pairs of taxa (here Anatolia vs. Asia, Anatolia vs. Africa, or Africa vs. Asia), and also synonymous polymorphisms (*P^S^*) and nonsynonymous polymorphisms (*P^N^*) within a taxon (here only for Africa or for Asia). We added 1 to the whole dataset to avoid zero division errors. [Note that we use the ratio between the numbers of substitutions, instead of estimating the rate of changes (i.e. *dN* and *dS*), because we are comparing ratios calculated for the same locus across the three lineages, and the denominators would cancel out]. We then calculated three statistics, *x*_1_, *x*_2_and *x*_3_ per gene, as follows:

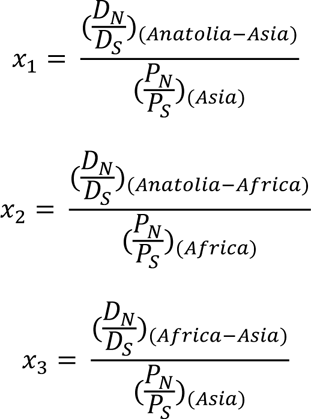

The *PMK* statistic was calculated as:

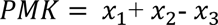

Here, *x*_1_, *x*_2_ and *x*_3_ values can be considered to measure the degree of non-synonymous differentiation of that gene between lineages. By summing Anatolia-Asia and Anatolia-Africa differentiation and subtracting Asia-Africa differentiation from this sum, we expect that the genes that have Anatolian-specific adaptations will appear as outliers with high PMK values. This is similar to the Population Branch Statistic (PBS), which uses the *F^ST^* statistic to identify loci differentiated in one population compared to two others (Yi et al., 2010).

After calculating all PMK scores for all genes, we calculated the *Z*-score of PMK values for each gene by scaling (subtracting the mean and dividing by the standard deviation) and considered genes with *Z*> 3 showing signs of putative positive selection. We applied Benjamini-Hochberg multiple testing correction on the *Z*-score-based p-values using the R *p.adjust* function.

### Non-synonymous mutation load

By using only transversions to avoid ancient damage, 2957 genes with *PAML*-estimated *dN/dS* values smaller than 1 were compared using paired Wilcoxon test of compare_means function of R ggpub package (version 0.4.0)(Kassambara, 2023). cdh008 representing hydruntine lineage did not show elevated *dN/dS* values compared to Asian and African lineages.

### Data and code availability

Genetic data generated from three Anatolian ancient equid samples were deposited to the European Nucleotide Archive ENA under project code PRJEB52847 as *BAM* files. The mitochondrial DNA sequence was deposited to the NCBI Genbank with accession no: OP448588.

The scripts incorporated into phylogenetic and demographic analyses were uploaded to https://github.com/CompEvoMetu/Hydruntine2023. The scripts related to selection analysis, namely, used for obtaining the polymorphism in the representative gene set, counting *D^N^*, *D^S^*, *P^N^* and *P^S^* and calculating PMK can be accessed at https://github.com/rabiafidan/equus.

## Supporting information

Archaeological Supplement

## Acknowledgements

The authors would like to thank Dr. Derya Baykara, for helpful comments; Dr. Mikolaj Lisowski and Dr. Jesse Wolfhagen for their contributions in sample selection; the Turkish Ministry of Culture and Tourism, Yozgat Museum and Konya Museum for permissions to work on archaeological samples; Sevgi Yorulmaz, Duygu Deniz Kazancı and Dr. Aslıhan Ilgaz for their assistance, and all METU CompEvo research team members for discussion and support.

This study was funded by a Scientific and Technological Research Council of Türkiye (TÜBİTAK) 1001 Grant Programme (Project No. 117Z991 to F.Ö.), the European Commission Horizon 2020 TWINNING Programme (Project No. 952317 “NEOMATRIX” to M.S.) and European Research Council Consolidator Grant H2020 ERC (no. 772390 “NEOGENE” to M.S.)

## Author Contributions

M.Ö., F.Ö. and M.S. conceived and designed the study and experiments. K.P., I.A., S.E.A., B.S.A., S.R.S. and G.M. prepared and provided zooarchaeological material. M.Ö. performed molecular biology laboratory experiments with support from A.A. and F.Ö.. M.Ö., K.G., E.Y., K.B.V., G.A., F.R.F., E.S., N.E.A. and D.K. analyzed data with the supervision of F.Ö., C.C.B., E.M.G., T.G., A.G., and M.S., Y.S.E., İ.T.. M.Ö., F.Ö. and M.S. wrote the manuscript with contributions from K.G., E.M.G., E.Y., G.A., F.R.F., E.S., I.H., B.S.A.. All authors read and approved the manuscript.

## Supplementary Tables

**Supplementary Table 1.** Pre-screening statistics of 15 equid samples. Samples which were sequenced further (see Supplementary Table 2) are indicated in bold. 5’ damage and 3’ damage indicate the proportion of C->T and G->A mismatches at first and last positions across all reads (Undet: Undetermined)

**Supplementary Table 2.** Deep sequencing statistics of selected individuals. Merged library (screening + deep sequencing) statistics are given in bold.

**Supplementary Table 3.** Partial mtDNA: samples for which only partial mtDNA sequences were available. Full mtDNA: samples for which full mtDNA sequences were available. Whole genome: samples for which whole or partial nuclear genome data were available.

**Supplementary Table 4.** f3 statistics for all possible comparisons based on autosomal chromosomes. (SE: standard error)

**Supplementary Table 5.** D-statistics for all possible comparisons on autosomal dataset. (SE: standard error)

**Supplementary Table 6.** D-statistics for all possible comparisons using the secondary autosomal variation dataset. (SE: standard Error)

**Supplementary Table 7.** List of genes tested and nominally significant (p < 0.05) in PAML codeml analysis.

**Supplementary Table 8.** List of genes tested, nominally significant (Z > 3) and significant after Benjamini-Hochberg multiple testing correction (p < 0.05) in Pairwise McDonald-Kreitman analysis. (BH: Benjamini-Hochberg)

**Supplementary Table 9.** Common genes that were both nominally significant in PAML codeml analysis (p < 0.05) and significant in PMK analysis after multiple testing correction (p < 0.05). Functional information was obtained from Ensembl Biomart Gene Ontology database.

