## Supplementary Figures for "The first complete genome of the extinct European wild ass (*Equus hemionus hydruntinus*)"

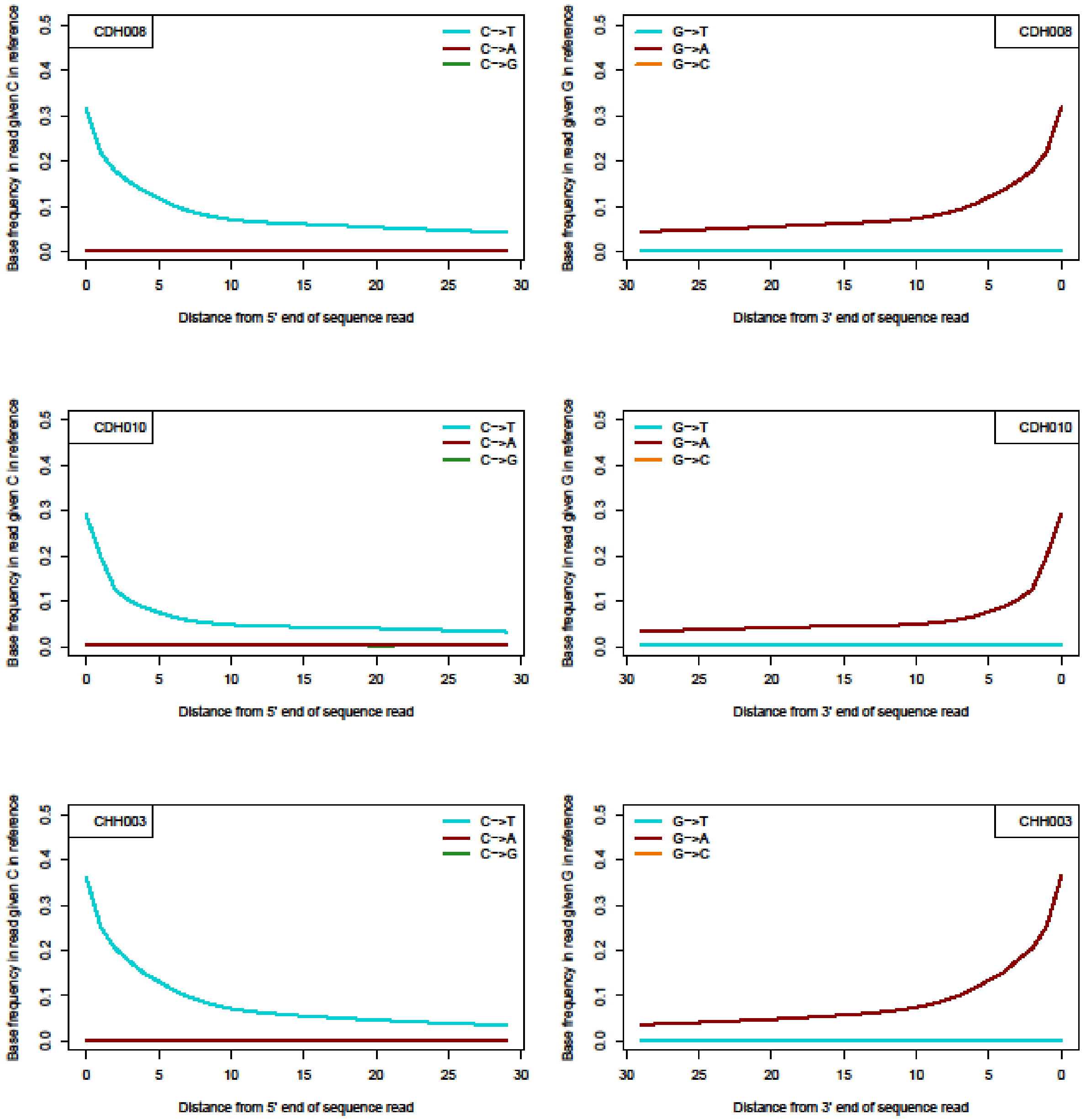


**Supplementary Figure 1.** Post-mortem damage signals of cdh008, cdh010 and chh003.

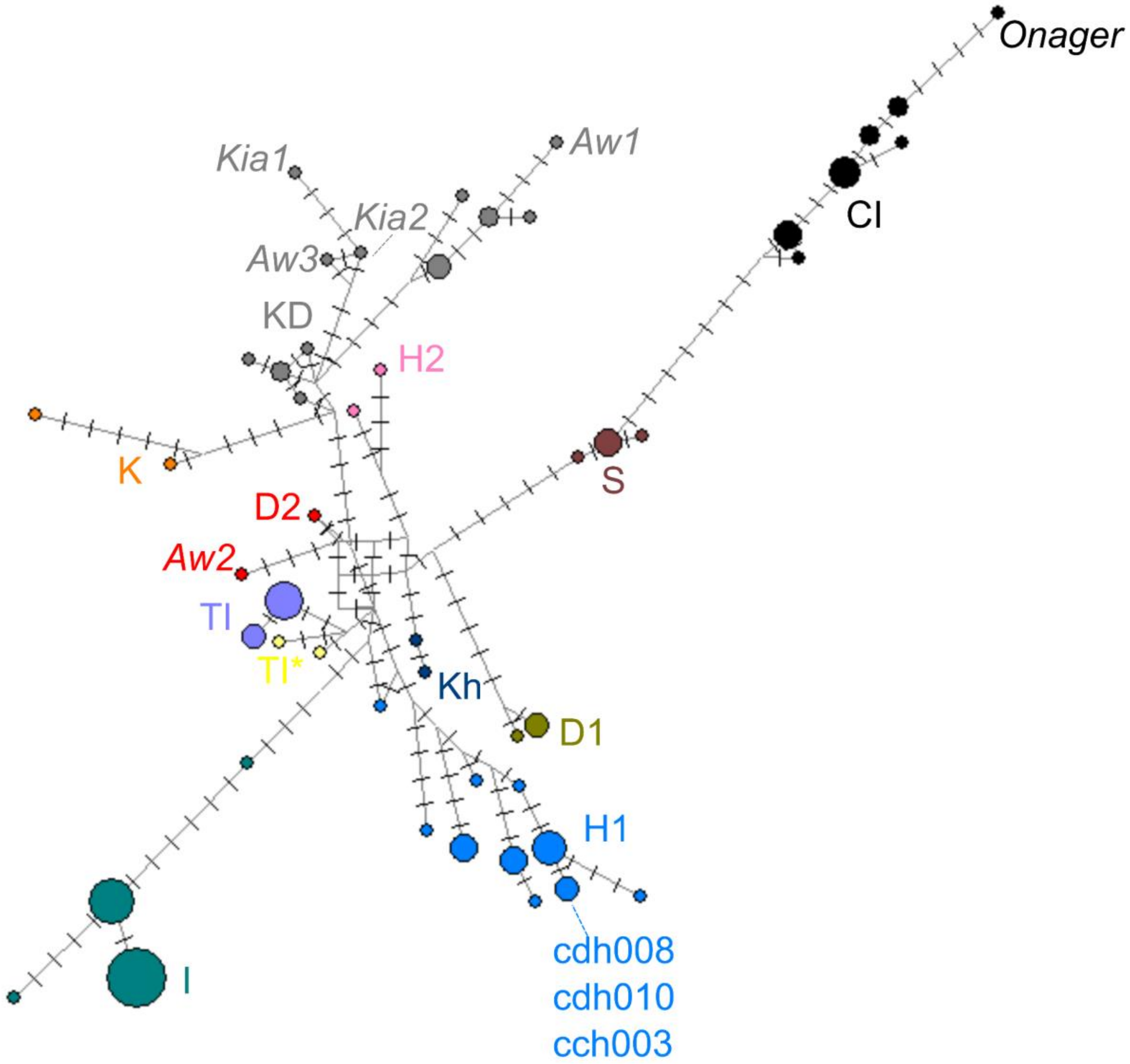


**Supplementary Figure 2.** Mitochondrial DNA median joining network of wild asses from Central/East Asia, Iran, India, Syria, Caucasus, Anatolia, Balkan and Western Europe (Orlando et al. 2006, Orlando et al.,2009, Bennett et al. 2017, Catalano et al. 2020), and sequences reported in this article generated using 249 bp diagnostic region of D-loop (Supplementary Tables 1 and 6, main text Methods section). (Refer to Supplementary Table 6 for the sample list, accession numbers and haplogroups. Also refer to methods about how samples were chosen. Names with italic font denote sample names and follow main text notation.)

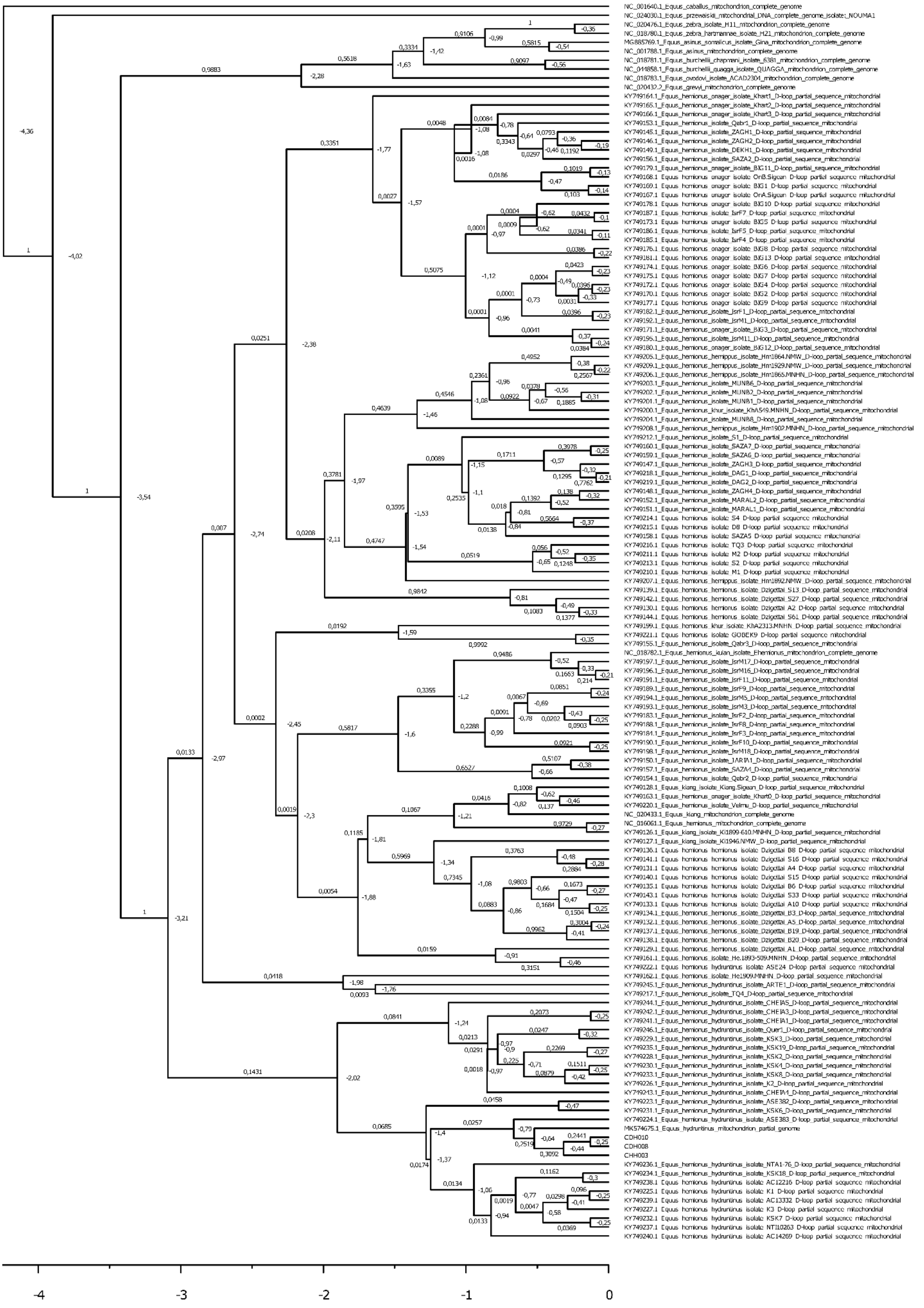


**Supplementary Figure 3.** The BEAST phylogenetic tree of 138 Equid samples generated using a 361 bp long partial sequence of the D-loop. In addition to the samples sequenced in this study and eight other Equid species, all samples used in Bennett et al. (2017) were included in the analyses given their availability in the corresponding region. *Equus caballus* was used as an outgroup, and a prior of 4.25 ± 0.125 millions of years (Mya) was given for the tree height, following Vilstrup et al. (2013). An uncorrelated relaxed clock and HKY nucleotide substitution model determined by  jmodeltest were used. The estimated divergence times are given in Mya on the nodes, and the posterior probability support values are on the branches [0, 1]. The reverse scale axis also shows time in Mya.

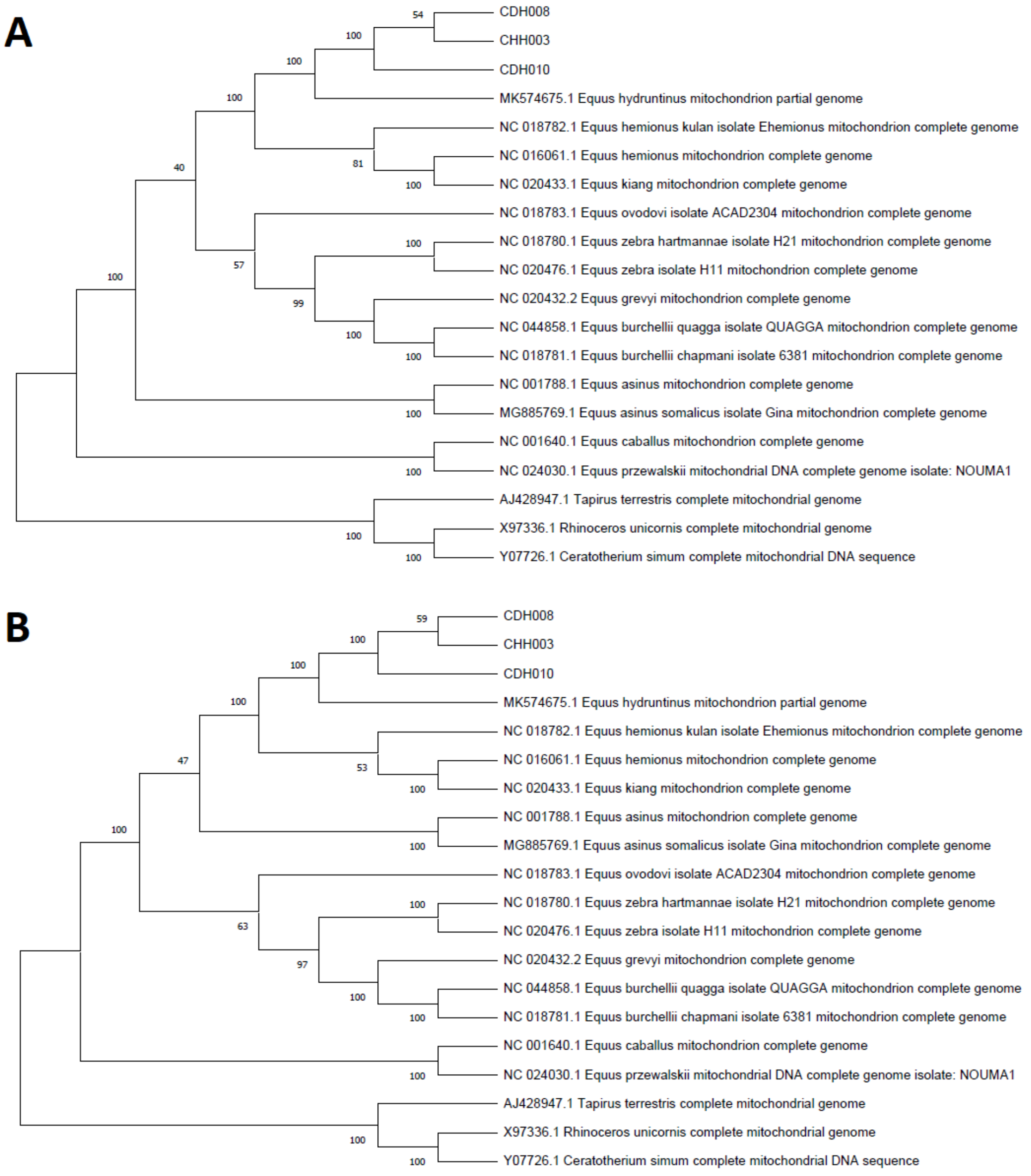


**Supplementary Figure 4**. A) Maximum likelihood (ML) and B) Neighbor-joining trees based on a 13,042 bp long consensus sequence of the mitochondrial genome. The nodal support was evaluated by 1000 bootstrap replicates for both methods. The substitution model of the ML tree was chosen as the GTR with Gamma distributed rate and invariant sites. The boostrap values are indicated on the nodes.

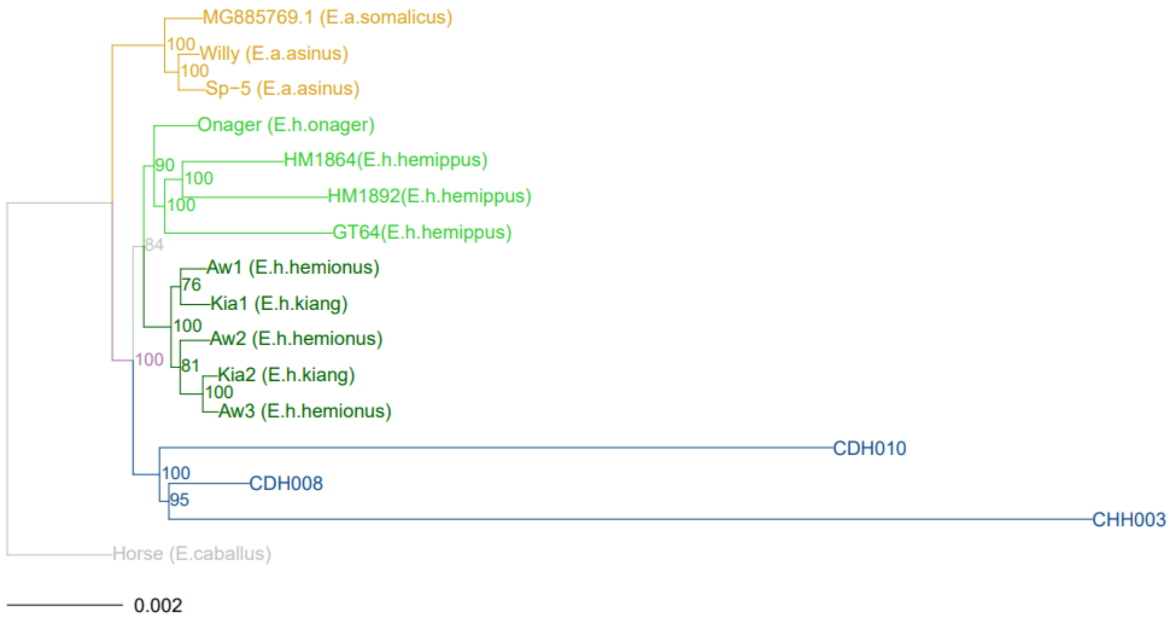


**Supplementary Figure 5.** A maximum likelihood tree constructed using concatenated protein coding sequences (29,384,180 bp) across the 15 genomes using only transversion sites. Numbers on the internal nodes show bootstrap support. External nodes indicate the genome/individual IDs, with species names in parentheses. Scale at the bottom shows branch lengths in terms of substitution per site.

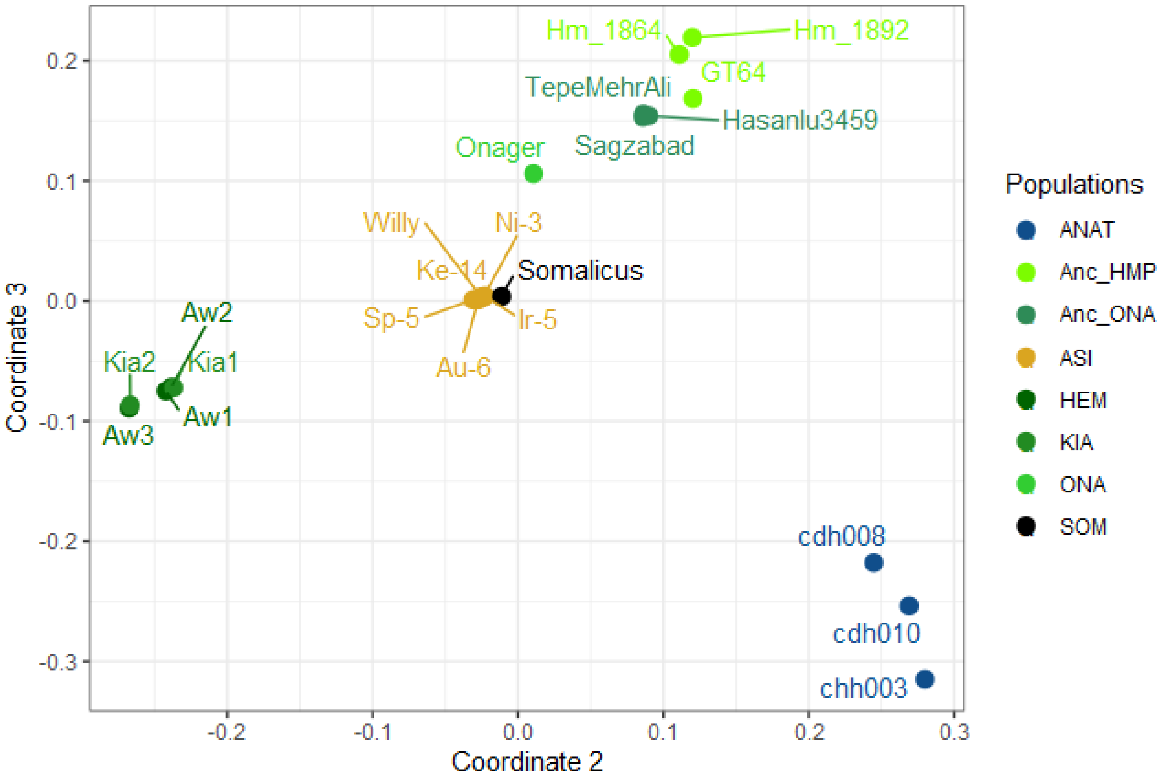


**Supplementary Figure 6.** Individual based multidimensional scaling plot on autosomal variation data (2,146,416 positions) showing the second and third coordinates. Color coding and labels are the same as Figure 3 in the main text. (ANAT: Anatolian samples reported in this article; Anc_HMP: Ancient *E.h.hemippus*; Anc_ONA: Ancient *E.h.onager*; ASI: *E.asinus*; HEM: *E.hemionus*; KIA: *E.h.kiang*; ONA: *E.h.onager*; SOM: *E.somalicus*)

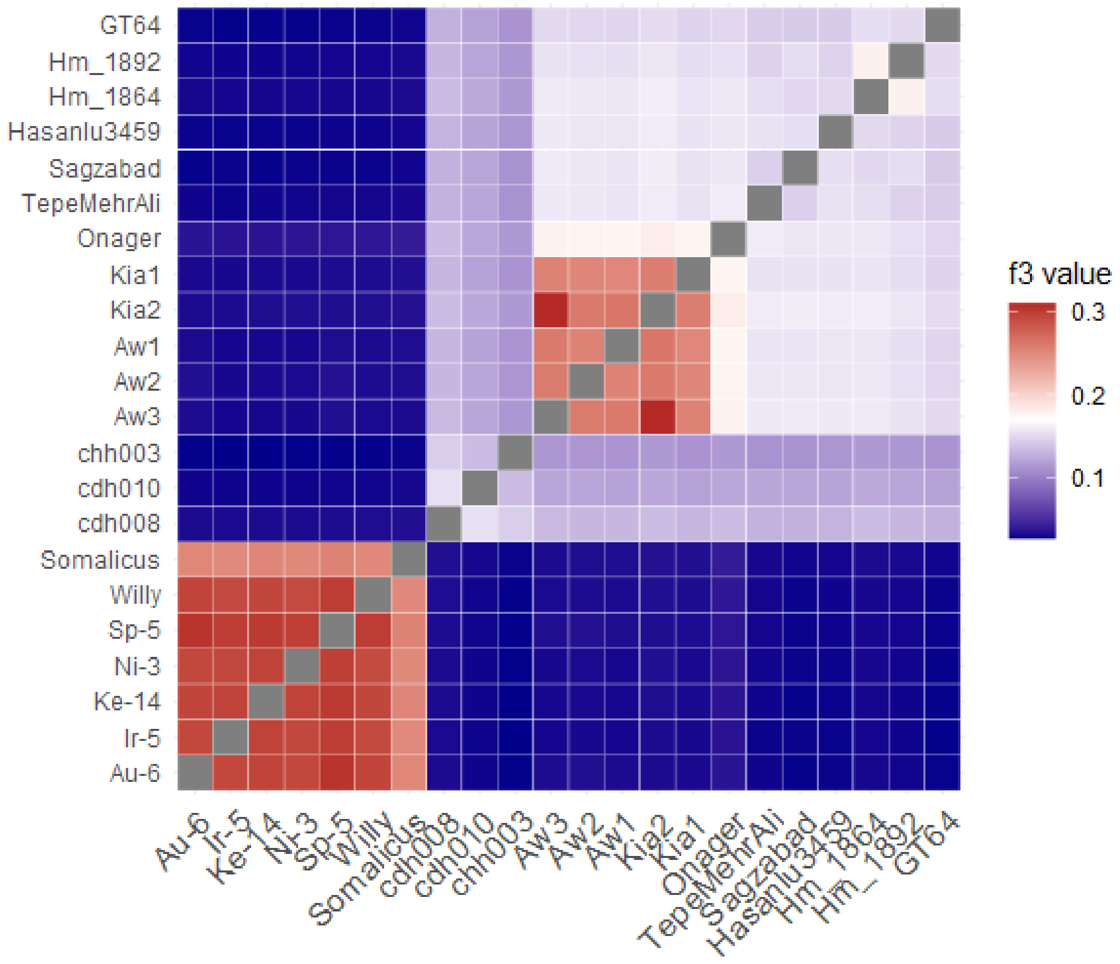


**Supplementary Figure 7.** Heatmap generated using outgroup-f3 data on autosomal loci total of 2,146,416 positions on individual level.

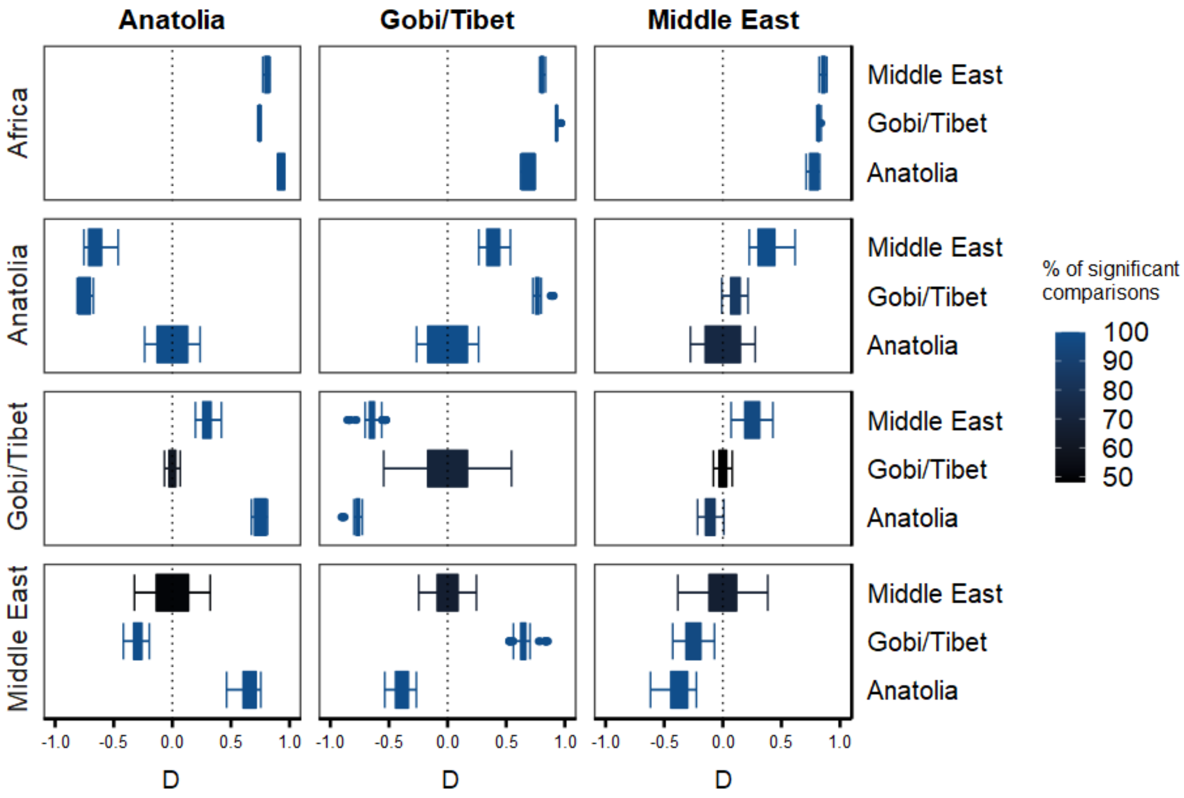


**Supplementary Figure 8**. D-statistics calculated using the secondary autosomal variation loci published previously (2,408,064 positions, see Materials and Methods), between regional groups. Test groups are shown on the top. Color gradient from blue to black represents the fraction of comparisons that are statistically significant (Z>3). Regional groups include: Anatolia (cdh008, cdh010, chh003), Gobi/Tibet (Aw1, Aw2, Aw3, Kia1, Kia2), Middle East (GT64, Hm_1864, Onager, Hasanlu3459, Sagzabad) and Africa (Asinus, Somalicus).

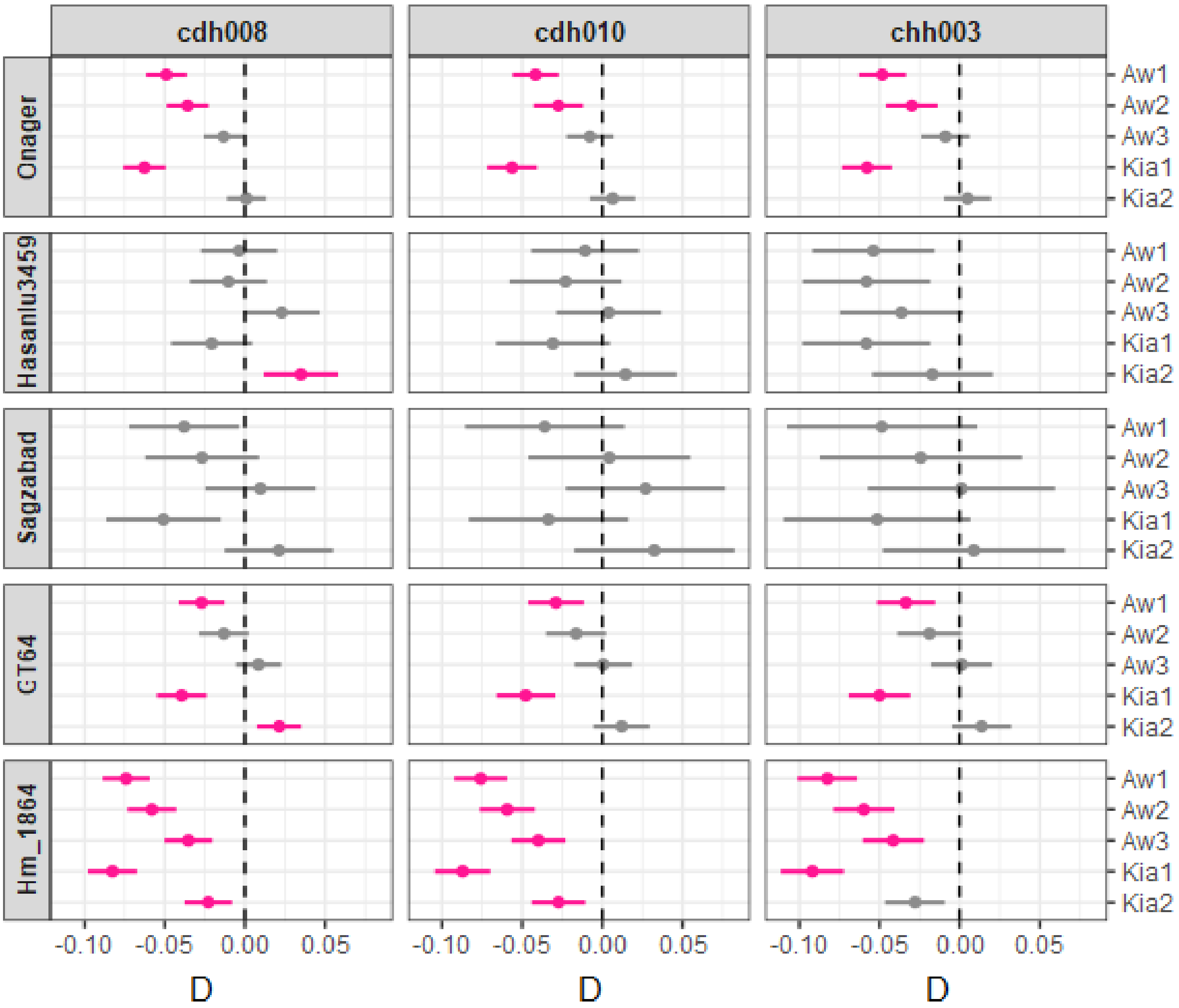


**Supplementary Figure 9.** D statistics results in the form of D(*Horse*,*Anatolia*; *MiddleEast*, *Gobi/Tibet*) based on the main autosomal dataset consisting of 2,146,461 variable sites. These are the same results as presented in summary form in main text Figure 5. Pink color represents significant results at Z>3 and gray color Z<3.

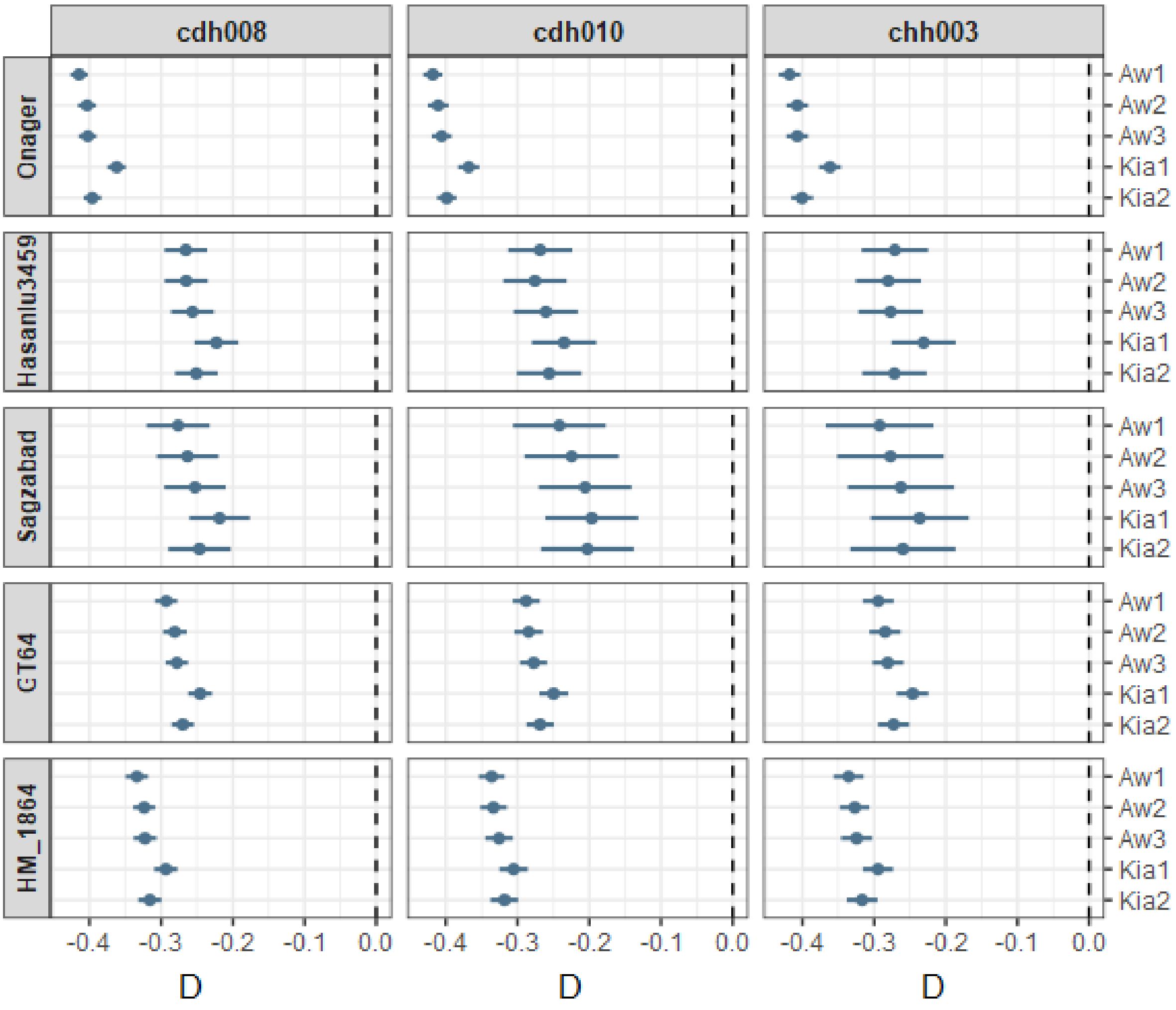


**Supplementary Figure 10.** D statistics results in the form of D(*Horse*,*Anatolia*; *MiddleEast*, *Gobi/Tibet*) based on the secondary autosomal variable loci (2,408,064 positions). Blue color represents significant results at Z>3 and gray color Z<3.

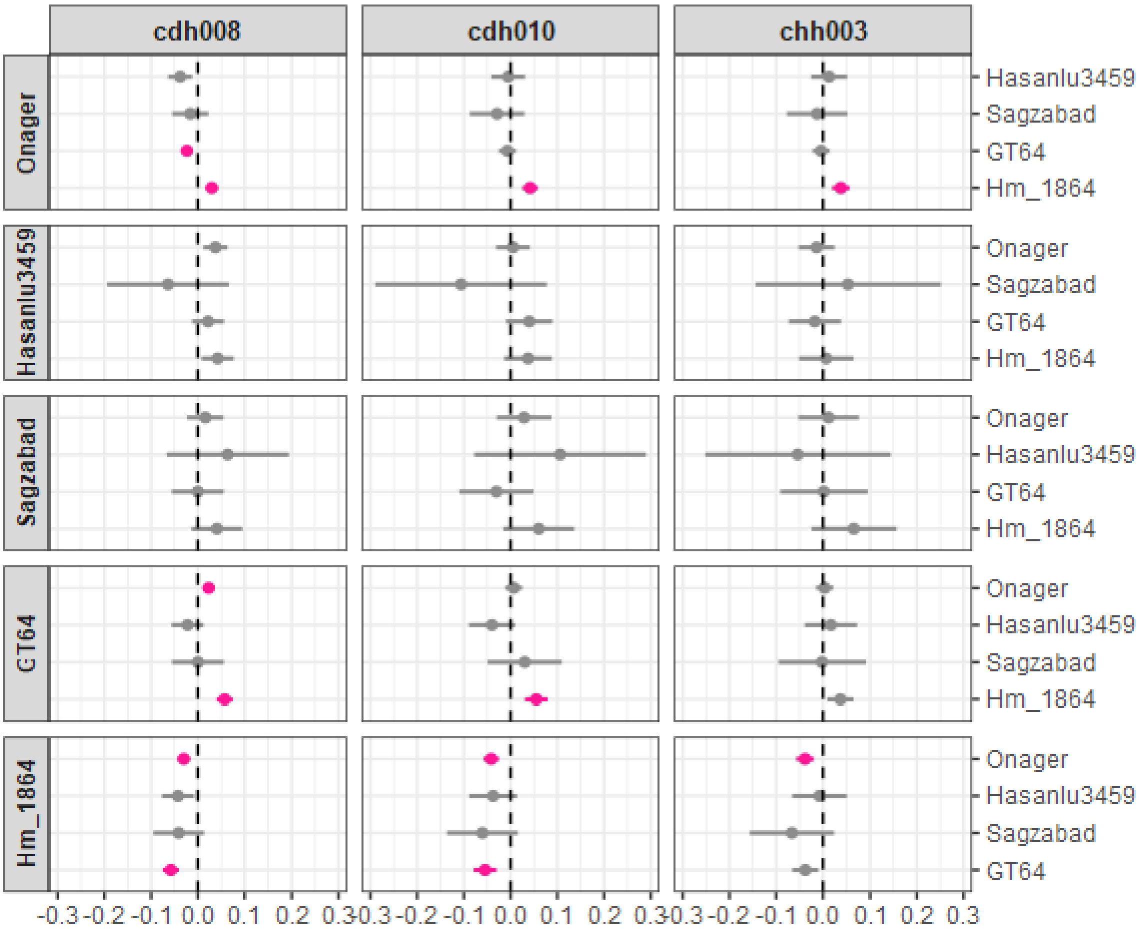


**Supplementary Figure 11.** D statistics results in the form of D(*Horse*,*Anatolia*; *MiddleEastX*, *MiddleEastY*) based on the main autosomal dataset consisting of 2,146,461 variable sites. These are the same results as presented in summary form in main text Figure 5. Pink color represents significant results at Z>3 and gray color Z<3.

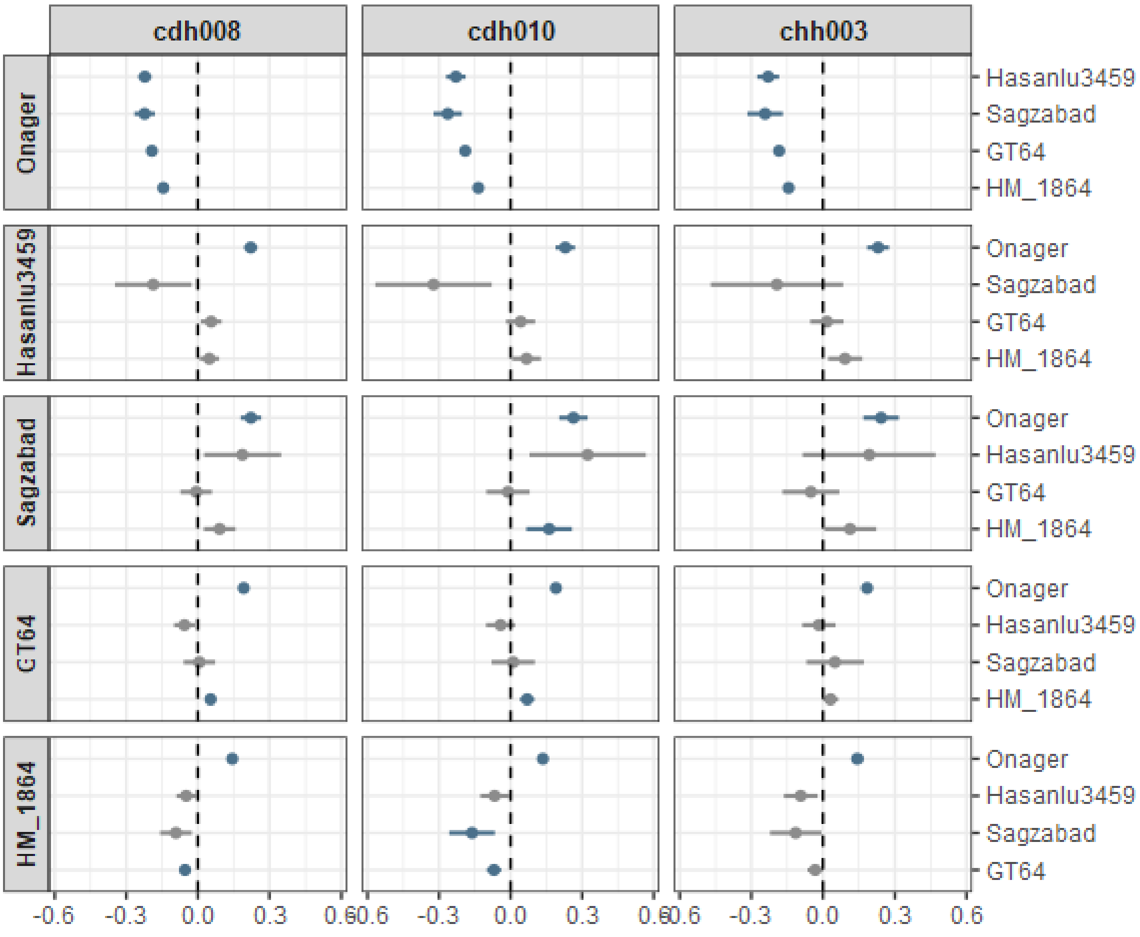


**Supplementary Figure 12.** D statistics results in the form of D(*Horse*,*Anatolia*; *MiddleEastX*, *MiddleEastY*) based on the secondary autosomal variating loci(2,408,064 positions). Blue color represents significant results at Z>3 and gray color Z<3.

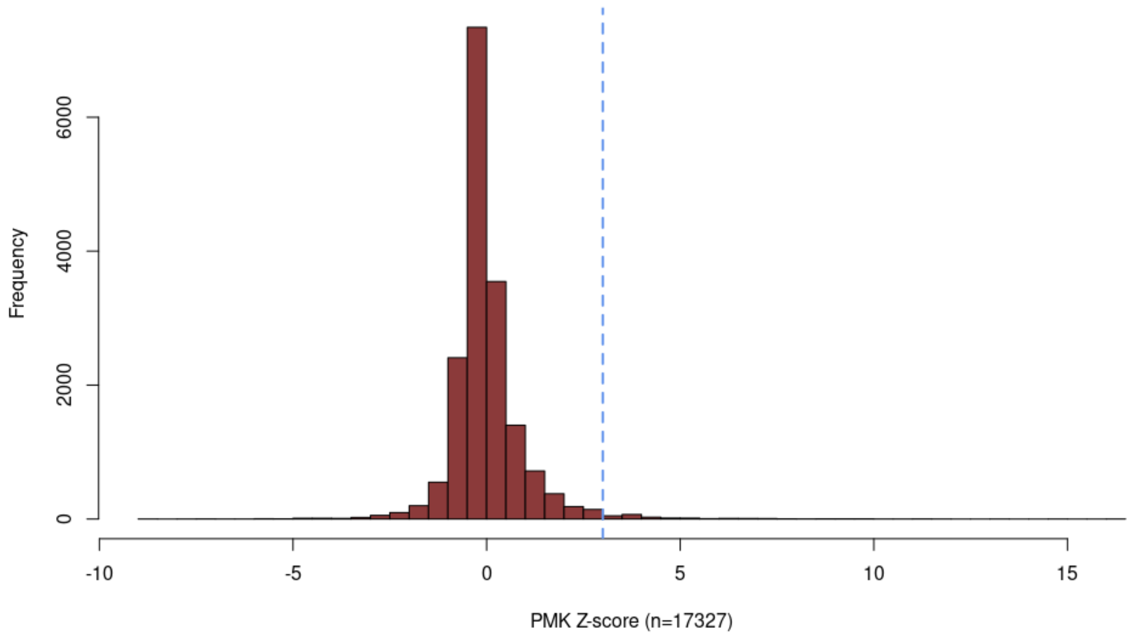


**Supplementary Figure 13.** Z-transformed Pairwise McDonald-Kreitman statistic (PMK) values of 17,327 protein-coding genes passing out filters. The blue vertical line marks Z=3.

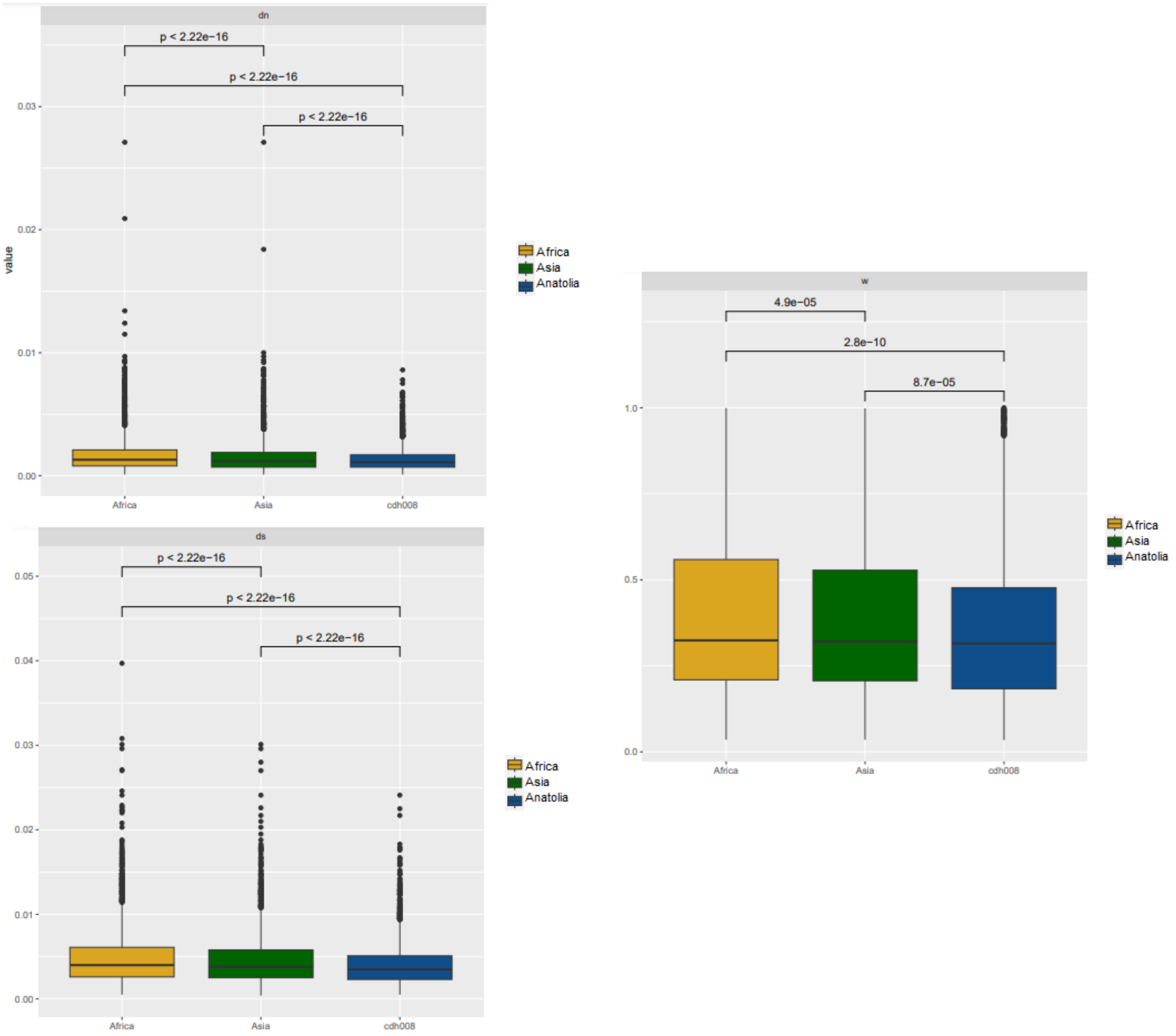


**Supplementary Figure 14.** dN, dS, and w values for African, Asian, and Anatolian lineages.
