## Supplementary material for "The first complete genome of the extinct European wild ass (*Equus hemionus hydruntinus*)": Archaeological Supplement

**Archaeological Supplement Table 1.** Bronze and Iron Age archaeological sites where Equids have been reported among faunal remains in modern-day Turkey. EBA: Early Bronze Age; MBA: Middle Bronze Age; LBA: Late Bronze Age; BA: Bronze Age; IA: Iron Age.

| Site | Period | Equus hydruntinus | Equus hemionus hemionus | Equus hemionus onager | Equus asinus asinus | Equus caballus | Equus sp. | Reference |
|---|---|---|---|---|---|---|---|---|
| Küllüoba | EBA | **?** | | | | | **X** | Efe and Fidan, 2008 |
| Acemhöyük | EBA | **X** | | | **X** | **X** | | Arbuckle, 2013 |
| Büyükkaya (Boğazköy) | EBA | | | | **X** | | **X** | von den Driesch and Pöllath, 2004 |
| Kaman Kalehöyük | EBA | | | | | | **X** | Atıcı, 2003 |
| Tilbeşar Höyük | EBA | | | **X** | | | **X** | Berthon and Mashkour, 2008 |
| Karataş Semayük | EBA | | | | | | **X** | Hesse and Perkins, 1974 |
| Büyüktepe | EBA | | **X** | | **X** | **X** | | Howell-Meurs, 2001b |
| İkiztepe | EBA | | | | | | **X** | Yakar, 2007 |
| Titriş Höyük | EBA | | | | | | **X** | Greenfield, 2002 |
| Gritille | EBA | | | | **X** | | | Stein, 1987 |
| Sos Höyük | EBA | | | | **X** | **X** | | Howell-Meurs, 2001a |
| Altıntepe | EBA | | | | **X** | **X** | **X** | Satar, 2006 |
| Çadır Höyük | MBA | | | | | | **X** | Adcock 2020 |
| Giricano | MBA | | | | | **X** | **X** | Berthon, 2013 |
| Kenan Tepe | MBA | | | | **X** | | **X** | Berthon, 2009 |
| Hibermerdon | MBA | | | | **X** | **X** | **X** | Berthon, 2009 |
| Başur Höyük | MBA | | | | | **X** | **X** | Berthon, 2013 |
| Ziyaret Tepe | MBA | | | | **X** | **X** | **X** | Greenfield - |

| Site | Period | Equus hydruntinus | Equus hemionus hemionus | Equus hemionus onager | Equus asinus asinus | Equus caballus | Equus sp. | Reference |
|---|---|---|---|---|---|---|---|---|
| | | | | | | | | Jongsma, 2013 |
| Kilisetepe | MBA | | | | | | X | Bursa, 2007 |
| Horum Höyük | MBA | | | | | | X | Bartosiewicz, 2005 |
| Kesikkaya Northwest (Boğazköy) | MBA | | | | | | X | Adcock, 2020 |
| Kültepe | MBA | | | | X | | | Atıcı, 2014b |
| Demirci Höyük | MBA | X | | | | | | Uerpmann, 1987 |
| Büyükkaya (Boğazköy) | LBA | | | | X | X | X | von den Driesch and Pöllath, 2004 |
| Boğazköy – Lower Town | LBA | | | X | X | X | X | von den Driesch and Boessneck, 1981 |
| Hibermedon | LBA | | | | X | X | X | Laneri et al, 2008 |
| Şarhöyük | LBA | | | | X | | | Satar, 2004 |
| Çadır Höyük | LBA | | | | | | X | Arbuckle, 2009; Adcock 2020 |
| Kavuşan Höyük | LBA | | X | | X | X | X | Berthon, 2013 |
| Giricano | LBA | | | | X | X | | Berthon, 2013 |
| Müslümantepe | LBA | | | | X | | X | Berthon, 2013 |
| Kilisetepe | LBA | | | | | | X | Gündem, 2015 |
| Sarıkale (Boğazköy) | LBA | | | | X | | X | Hollstein and Middea 2014 |
| Çatalhöyük | BA | X | | | | X | X | Pawłowska, in press; Russell et al., 2003, 2004; Twiss etal., 2005 |
| Çatalhöyük | IA | | | | | | X | Pawłowska, |

| Site | Period | Equus hydruntinus | Equus hemionus hemionus | Equus hemionus onager | Equus asinus asinus | Equus caballus | Equus sp. | Reference |
|---|---|---|---|---|---|---|---|---|
| | | | | | | | | in press; Russell et al., 2003, 2004; Twiss etal., 2005 |
| Panaztepe | IA | | | | | | **X** | Tekkaya, 1992 |
| Sos Höyük | IA | **X?** | | | **X** | **X** | | Sagona, 1998 |
| Büyükkaya (Boğazköy) | IA | | | | **X** | **X** | **X** | von den Driesch and Pöllath, 2004 |
| Büyüktepe | IA | | **X** | | **X** | **X** | | Howell - Meurs, 2001b |
| Çadır Höyük | IA | | | | | | **X** | Arbuckle, 2009; Adcock 2020 |
| Gordion | IA | | | | | | **X** | Miller, 2009 |
| Kaman Kalehöyük | IA | | | | | | **X** | Hongo, 1997 |
| Kilisetepe | IA | | | | **X** | **X** | | Baker, 2008 |
| Maşat Höyük | IA | | | | **X** | **X** | **X** | Dönmez, 2010 |
| Kavuşan Höyük | IA | | **X** | | **X** | **X** | **X** | Berthon, 2013 |
| Hibermedon | IA | | **X** | **X** | **X** | **X** | **X** | Laneri, 2008 |
| Büyükardıç | IA | | | | | **X** | | Açıkkol and Yılmaz, 2005 |
| Altıntepe | IA | | | | **X** | **X** | **X** | Satar, 2006 |
| Boğazköy | IA | | | | **X** | **X** | | Deniz, 2015 |
| Kesikkaya South (Boğazköy) | IA | | | | | | **X** | Adcock 2020 |
| Korucutepe | | | | | **X** | **X** | | Boessneck and Von Den Driesch, 1974 |

**Archaeological Supplement Table 2.** Archaeological information of equid samples subjected in this study. (NA: not available).

| **aDNA Lab ID** | **Site** | **Excavation ID** | **Tissue** | **Tissue subtype** | **Uncalibrated C14 date (BP)** |
|---|---|---|---|---|---|
| cdh008 | Çadırhöyük | 24515 | Bone | Phalanx | 2801 ± 47 (TÜBİTAK-1683) |
| cdh010 | Çadırhöyük | 23073 | Bone | Calcaneus | 2820 ± 28 (TÜBİTAK-1682) |
| cdh013 | Çadırhöyük | 24957 | Bone | Phalanx | NA |
| cdh022 | Çadırhöyük | 6683 | Tooth | Lower tooth | NA |
| chh001 | Çatalhöyük | 21075 | Tooth | Lower tooth | NA |
| chh002 | Çatalhöyük | 16247.F5 | Tooth | Lower tooth | NA |
| chh003 | Çatalhöyük | 21003.F38 | Tooth | Lower tooth | 2435 ± 28 (TÜBİTAK-1684) |
| chh004 | Çatalhöyük | 19347.F3 F20 | Tooth | Lower tooth | NA |
| chh005 | Çatalhöyük | 17097.F5 F6 | Tooth | Lower tooth | NA |
| chh006 | Çatalhöyük | 19101.F22 | Tooth | Lower tooth | NA |
| chh007 | Çatalhöyük | 19216.F268 | Tooth | Lower tooth | NA |
| chh008 | Çatalhöyük | 15743.F1676 | Tooth | Lower tooth | NA |
| chh009 | Çatalhöyük | 20465.F115 | Tooth | Lower tooth | NA |
| chh0010 | Çatalhöyük | 30745.F21 | Tooth | Lower tooth | NA |
| chh0011 | Çatalhöyük | 19349.F7 F8 | Tooth | Lower tooth | NA |

### Archaeological Information on Excavation Sites

**Çatalhöyük**

Çatalhöyük is a tell site near Konya in central Turkey. First excavated by James Mellaart in the 1960s, a new project began in 1993 under the direction of Ian Hodder. At first the focus was on surface study and survey, followed by excavation from 1995 to 2017. The site is of international significance because it reached a large size (13.5 ha) at an early date, had a dense population (1,500 to 5,000 people), has rich symbolism and subfloor burials, and was occupied for 1,500 years; the Neolithic East mound dates from 7100 BCE to 5900 BCE with the Chalcolithic West Mound overlapping in time in the last quarter of the seventh millennium BCE and continuing on until 5600 BCE. The well-preserved buildings and rich art in the Neolithic mound give a unique insight into early village life. The site allows study of many of the main questions dealing with the early formation of settled villages/towns and the early intensification of agriculture. The site was inscribed on the UNESCO World Heritage list in 2012.

**Çadır Höyük**

ÇadırHöyük is located in the Yozgat Province in north central Turkey, near the modern city of Sorgun. The mound rises 32 m above the floodplain and covers an area of approximately 5 hectares, with an additional occupation area north of the mound. Investigations have been ongoing at Çadır since 1994. A 2 x 2 m sounding on the mound's lower southern slope documented that occupation at Çadır extends back to at least the Middle Chalcolithic (Beta 146707, 5220–4940 BC [Cal BP 7170–6890]). The mound has been continuously occupied since that time, with final abandonment in the 14$^{th}$ century CE. At present, a large expanse of horizontal exposure (ca. 700 m2) on the southern slope offers data on the Late Chalcolithic occupation of the mound (Steadman et al. 2018, 2019a, 2019c).

The Byzantine occupation is found in the area north of the mound (known at the North Terrace) which features a large Byzantine residential community (Cassis et al. 2019;Cassis and Steadman 2014; Steadman et al. 2019b). On the mound's summit there is an extensive Byzantine defensive wall, inside of which were storage areas; shortly before abandoning the community, residents moved up onto the summit for protection during a turbulent time (Cassis et al. 2019).Bronze Age occupation, including the Hittite Empire period, is found on the mound's eastern and northern slopes. A Middle Bronze wall, followed by a significant Hittite-era defensive wall ringed the mound in the Late Bronze Age (Ross et al. 2019b; Steadman et al. 2019b; Steadman and McMahon 2017). Inside the wall are residential areas and a small portion of a substantial Hittite structure. At present the Iron Age and Byzantine overburden covers much of the Bronze Age occupation on the mound.

The Iron Age occupation extends across much of the mound, but is currently largely covered by the Byzantine settlement. Excavation has identified a Late Iron staired pathway up the mound toward the summit, and a collection of Late Iron rooms, gates, and pathways into the summit occupation area (Steadman et al. 2019b; Steadman and McMahon 2017). It is likely that a significant Early–Late Iron Age occupation of the mound summit will be revealed through further excavation. The USS 4 trench, from which the four samples came, is located on the upper southern slope. Rather than residential, this area was likely outside the main occupation area and was devoted to craft production including textiles, followed by what appears to be felt and leather production (Ross et al. 2019a, 2019b). The CDH08 (USS4 F128, FCN11128) and CDH010 (USS4 L198, FCN11130) samples derive from a stone wall (F128) set into a subfloor (L198) that we have phased to the Middle Iron Age. The CDH013 sample comes from a matrix phased to the transition between the Early and Middle Iron Age. The CDH022 (USS 4 L281, FCN 14393) sample comes from the fill in a pit (F212) that rests on the cusp between the very end of the Late Bronze Age and the beginning of the Early Iron Age.

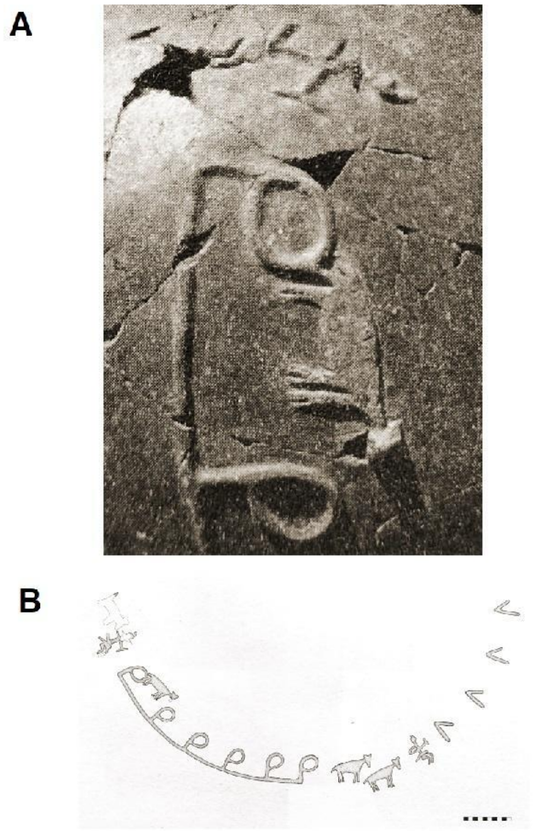

**Archaeological Supplement Figure 1.** Pottery fragment with decorations of a hunting scene of equids (possibly a wild ass) from Neolithic Köşk Höyük in Central Anatolia (credit: Aliye Öztan).

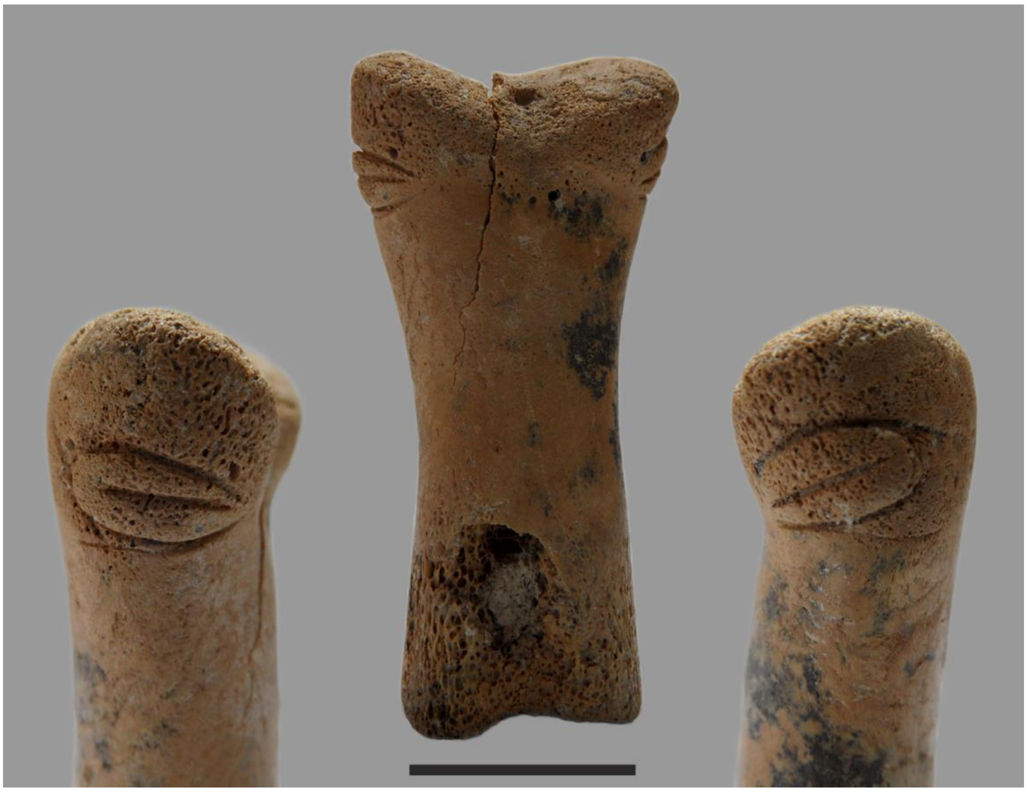

**Archaeological Suplement Figure 2.** An equid bone figure from Çatalhöyük (Photo by J. Quinlan).
